# Direct photoreception of a pituitary endocrine cell, melanotroph, induces a hormone release

**DOI:** 10.1101/2023.08.02.551597

**Authors:** Ayaka Fukuda, Keita Sato, Chika Fujimori, Takahiro Yamashita, Atsuko Takeuchi, Hideyo Ohuchi, Chie Umatani, Shinji Kanda

## Abstract

In addition to canonical photoreception by the eye, many other organs express non-visual photoreceptors although their biological significance is mostly unknown. Here, we discovered a novel phenomenon in which the pituitary of medaka directly receives light, which induces hormone release. Ca^2+^ imaging analysis revealed that a melanotroph, a pituitary endocrine cell secreting melanocyte-stimulating hormone (MSH), robustly increases [Ca^2+^]_i_ during short-wavelength light irradiation. Moreover, we identified Opn5m as the key molecule of this mechanism. The significance of this phenomenon was suggested to be involved in UV protection because knockout of *opn5m* significantly reduced the expression of *tyrosinase*, the rate-limiting enzyme for melanogenesis, in the skin. These results suggest a novel mechanism in which direct reception of short-wavelength light by pituitary endocrine cells triggers the pathway to enhance UV protection.

**One-Sentence Summary:** An endocrine cell of the pituitary was proven to be a photoreceptive cell that enables autonomous hormone release.

## Introduction

A growing body of evidence shows that various organs, including the central nervous system, express non-visual opsin genes that may contribute to the perception of the light information in vertebrates (*1, 2*). However, very few phenomena have had their significance and mechanisms explained at the molecular and cellular level. For instance, the pineal gland in non-mammalian vertebrates is known to possess photoreceptive cells and photoreception by the pineal gland modulates circadian rhythms (*3, 4*). This also involves the interesting evolutionary event in which this direct photoreception mechanism has been lost in mammals, which generally have untransparent head structures. In these species, the modulation of circadian rhythm by light is exclusively achieved by the light information from the eye via a neural pathway (*5*). Considering this fact, non-mammalian models with relatively transparent skulls may exhibit intriguing non-visual phenomena. Here, using a non-mammalian model teleost, medaka, which is amenable to the application of various technologies (*6-9*), we found a surprising pathway suggesting that melanotroph, an endocrine cell in the pituitary, which is located at the base of the brain, directly receive the light and autonomously secrete the hormones. The present study aimed to investigate the mechanism and biological significance of this phenomenon.

## Results

First, we performed Ca^2+^ imaging of melanotrophs using a whole brain-pituitary *in vitro* preparation of *pro-opiomelanocortin* (*pomc*):GCaMP medaka. Under the *cis-*regulatory activity of the 3.7kb 5’ flanking region of *pomc*, melanotrophs and corticotrophs were specifically labeled by a Ca^2+^ indicator, GCaMP6s (*10*) (fig. S1). During Ca^2+^ imaging, a robust [Ca^2+^]_i_ increase in melanotrophs was observed without any additional stimulation except the exposure to the blue excitation light for fluorescence observation. This phenomenon suggests that some region in the brain or pituitary received the light and induced the [Ca^2+^]_i_ increase in the melanotrophs (fig. S2). To determine whether the light is sensed by the pituitary or brain, we performed Ca^2+^ imaging using an isolated pituitary. It was indicated that, even in this isolated pituitary, the [Ca^2+^]_i_ increase is observed in melanotrophs within a few seconds in response to the blue excitation light exposure (Fig. 1A). Note that corticotrophs, which express adrenocorticotropic hormone (ACTH) (*11*) derived from the same precursor, POMC, did not exhibit the [Ca^2+^]_i_ increase under the same condition (Fig. 1B). Since this robust rise in [Ca^2+^]_i_ triggers hormone release in endocrine cells (*12-14*), it was strongly suggested that isolated pituitary can autonomously release MSH and/or its derivatives by directly sensing light. Next, we analyzed the wavelength specificity of this non-visual photoreception of the pituitary using stimulation light with different wavelengths. To avoid the unintentional direct detection of stimulation light during imaging, stimulation light was irradiated only during the interval of each imaging acquisition by using an Arduino (microcontroller)-based control system (fig. S3). In this experiment, the effects of light stimulations (365–740 nm, approximately 200 μmol m^-2^ s^-1^) during recording intervals were evaluated (excitation exposure = 50 ms, recording interval = 5 s). Because it was proven that melanotrophs show a steady [Ca^2+^]_i_ level after 200 s under this imaging condition (fig. S4A), we started the evaluation at least 200 s after the 5 s interval fluorescence image acquisition started. The time course of this experiment is detailed in fig. S4B. Here, we identified that melanotrophs respond strongly to short-wavelength light, especially UV light (Fig. 1C and D), which suggests that a short-wavelength sensitive photoreceptor is involved in this photoreception mechanism. Next, to examine whether a melanotroph directly receive the light or if another photoreceptive cell in the pituitary mediates this light-induced [Ca^2+^]_i_ increase, we performed Ca^2+^ imaging of dissociated cells from the posterior pituitary of *pomc*:GCaMP medaka. It was demonstrated that an isolated melanotroph labeled by GCaMP responds to the excitation light (Fig. 1E), which is similar to the response observed in the whole pituitary preparation. As expected, three repetitive trials caused [Ca^2+^]_i_ increase of an isolated cell (fig. S5). These results demonstrate that a melanotroph, an endocrine cell, receives short-wavelength light and increases the [Ca^2+^]_i_. Given that melanotrophs are endocrine cells, this suggests a novel mechanism involving the autonomous release of hormone in response to light. Although such hormone release from endocrine cells directly induced by light has not yet been reported, these data clearly indicate the existence of this novel mechanism. Additionally, we explored the Ca^2+^ sources of the extracellular or intracellular stores responsible for this light-induced [Ca^2+^]_i_ increase. Perfusion experiments using modified artificial cerebrospinal fluid (ACSF) or normal ACSF containing inhibitors showed that Ca^2+^-free ACSF or an inhibitor of the Ca^2+^ channel, Cd^2+^ (CdCl_2_), did not affect the light-induced [Ca^2+^]_i_ increase, whereas an inhibitor of Ca^2+^ release from endoplasmic reticulum (ER), 2-Aminoethoxydiphenyl Borate (2-APB), drastically diminished the light-induced [Ca^2+^]_i_ increase (Fig. 1F and fig. S6). Consequently, it is suggested that Ca^2+^ can be derived from intracellular ER.

**Fig. 1.**
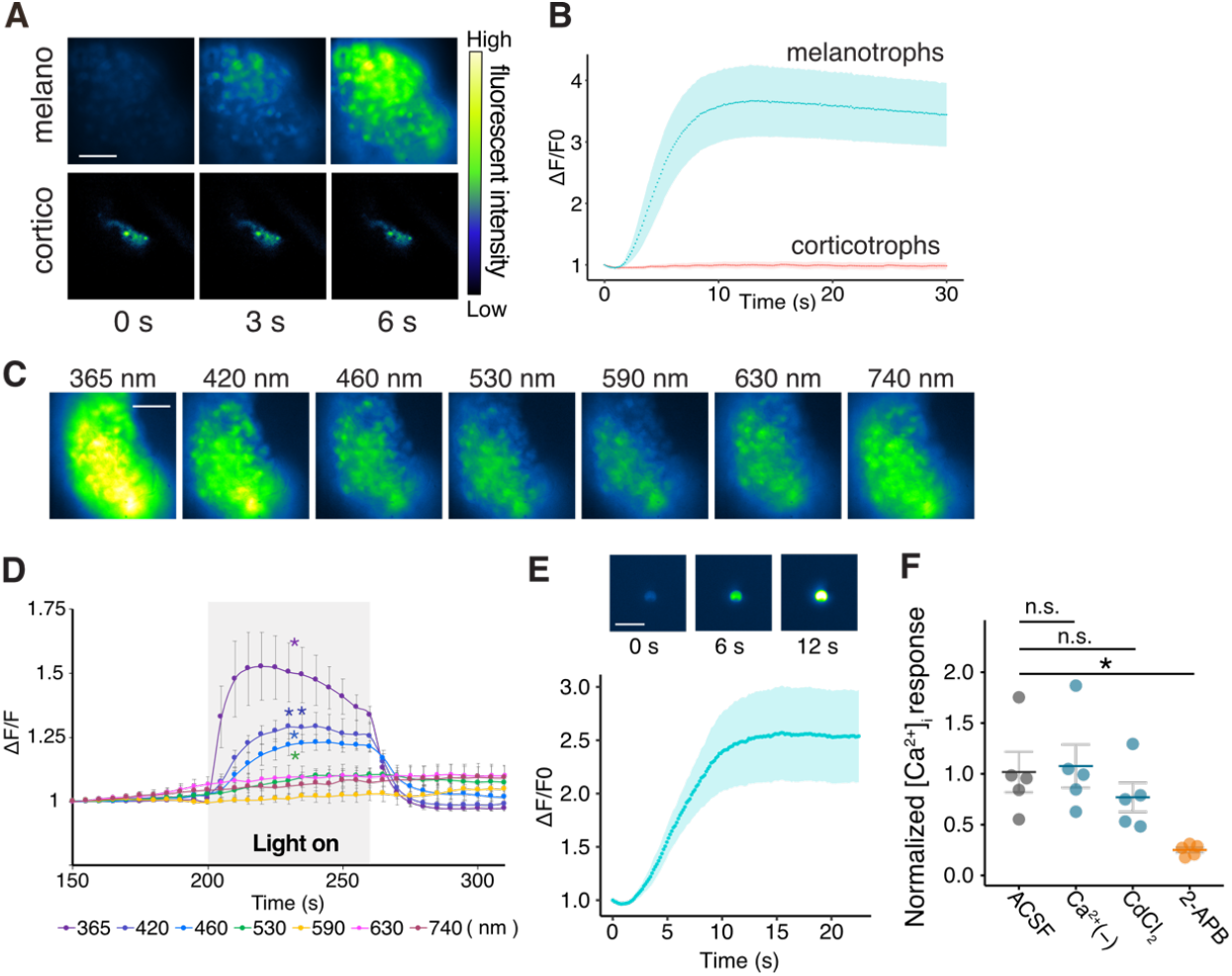
Short-wavelength light induced [Ca^2+^]_i_ rise of melanotrophs in the pituitary. **(A)** Representative image series showing fluorescence changes during the excitation light exposure (450–490 nm) in melanotrophs (melano) and corticotrophs (cortico) of *pomc*:GCaMP medaka. **(B)** Time course of GCaMP fluorescence changes of melanotrophs and corticotrophs. *N* = 5 medaka. **(C)** Representative GCaMP fluorescence images of melanotrophs 20 s after each wavelength (365, 420, 460, 530, 590, 630, 740 nm) light irradiation. Note that all irradiated light was adjusted to 200 (μmol m^-2^ s^-1^) photon flux density (PFD). **(D)** Time course of GCaMP fluorescence changes of melanotrophs with various wavelength light stimulation. Each wavelength light was irradiated 200 to 260 s after the start of recording. *n* = 5 medaka, paired *t* tests were applied between before and during irradiation for each wavelength. **(E)** Representative images of an isolated melanotroph labeled with GCaMP showed fluorescence increase during imaging with the blue excitation light (450–490 nm). Time course of GCaMP fluorescence changes of melanotrophs and corticotrophs. *n* = 5. **(F)** Ca^2+^ response to the blue excitation light in melanotrophs during pharmacological inhibition using Ca^2+^-free ACSF, 100 μM CdCl_2_, and 500 μM 2-APB. *n* = 5 medaka; *P* = 0.011; n.s., not significant; Dunnet tests. Scale bar, (A, C) 50 μm and (E) 25 μm. Data are represented as mean ± SEM. \**P* < 0.05, \*\**P* < 0.01, \*\*\**P* < 0.001.

Next, we identified the photoreceptor protein involved in this direct photoreception in melanotrophs. Given the results of *in situ* hybridization, which showed *opn5m* is expressed in the posterior part of the pituitary (fig. S7) (*15*), we performed double *in situ* hybridization in conjunction with *pomc* (Fig. 2A). We found co-expression of *opn5m* and the *pomc* gene in the posterior part of the pituitary, which indicates that melanotrophs express *opn5m* mRNA. Subsequently, recombinant Opn5m was prepared, and the UV-visible absorbance spectrum was analyzed. The 11-*cis*-retinal-bound resting state of Opn5m showed the absorption maximum at the short-wavelength region (Fig. 2B and fig. S8), which is consistent with the property of zebrafish Opn5m (*16, 17*). These results are in agreement with those of Ca^2+^ imaging showing that melanotrophs responded strongly to short-wavelength light (Fig. 1D). Next, to examine the possibility that Opn5m is the key molecule of the light sensitivity of melanotrophs, we disrupted the *opn5m* gene of *pomc*:GCaMP medaka using CRISPR/Cas9 (*18*) (fig. S9) and performed Ca^2+^ imaging. Here, *opn5m*^-/-^ medaka did not respond to the excitation light, whereas *opn5m*^+/-^ responded as observed in the wild type (WT) (Fig. 2C to E). Thus, it was proven that Opn5m is exclusively important for the short-wavelength light-induced [Ca^2+^]_i_ increase of melanotrophs. To confirm that this defect is specific to light sensitivity, we examined the effect of corticotropin-releasing hormone (CRH), which is known to induce the release of MSH in another teleost (*19*). Note that the receptor of CRH, *crhr1*, was suggested to be expressed in melanotrophs in medaka as well (fig. S10). Unlike the *opn5m*-dependent mechanism of light stimulation, CRH increased the [Ca^2+^]_i_ in both *opn5m*^+/-^ and *opn5m*^-/-^ individuals (Fig. 2F and fig. S11). Furthermore, we clarified whether Opn5m is sufficient for the light-induced [Ca^2+^]_i_ increase by conducting Ca^2+^ imaging of a widely used human cell line, HEK293A cells, or a rodent pituitary cell line, LβT2 cells (*20*), transfected with *opn5m* and *gcamp6s*. As expected, after incubation with an opsin chromophore, 11-*cis*-retinal or all-*trans*-retinal, these HEK cells and LβT2 cells also showed a [Ca^2+^]_i_ increase while the blue excitation light was irradiated (Fig. 2G, H and fig. S12), which is consistent with previous studies of mammalian Opn5m (*21-23*). From these lines of evidence, we conclude that Opn5m is necessary and sufficient for the [Ca^2+^]_i_ increase caused by photoreception.

**Fig. 2.**
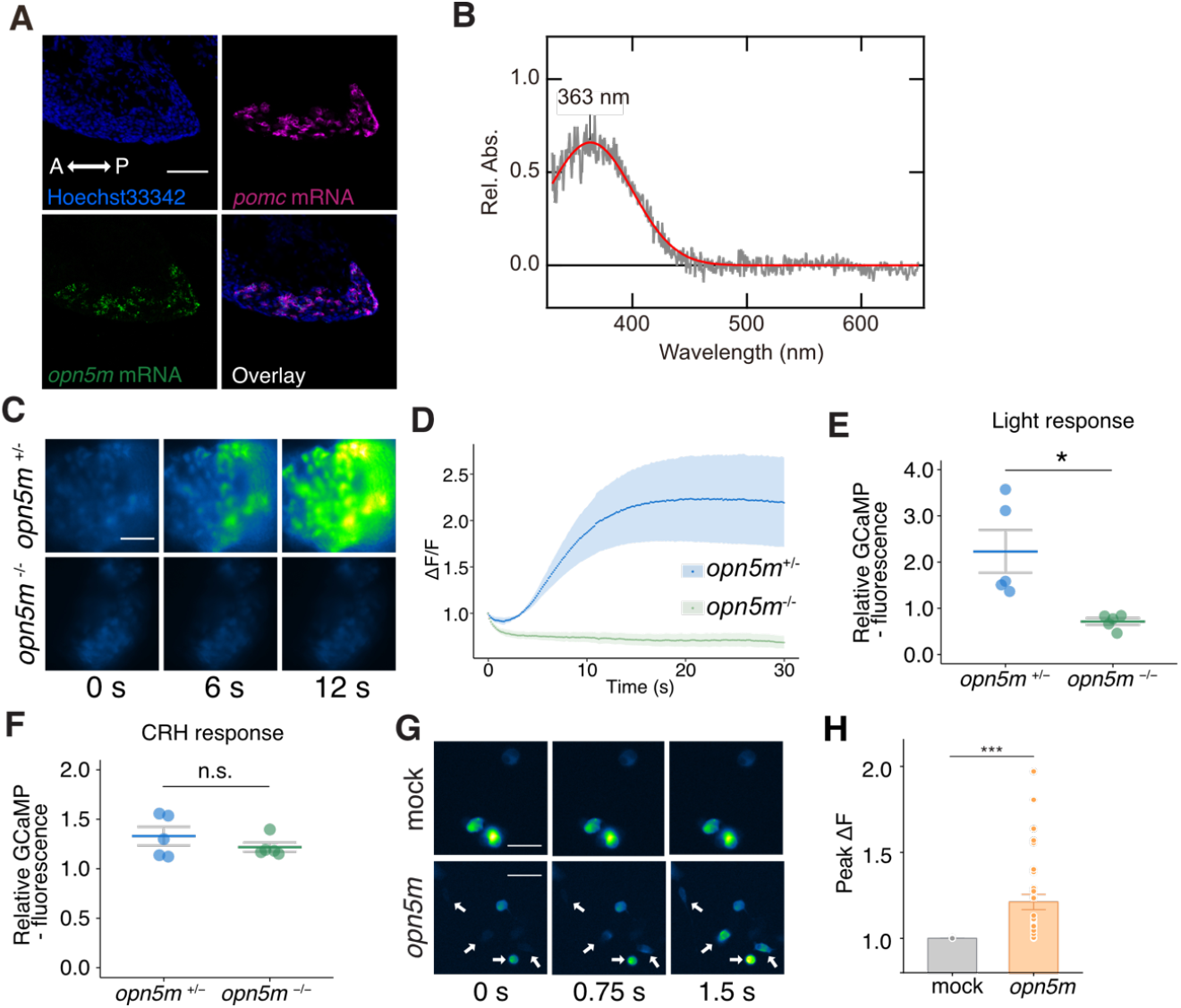
Opn5m is necessary and sufficient for light sensitivity of melanotrophs. **(A)** Double *in situ* hybridization indicates that *opn5m* is expressed in melanotrophs. Scale bar, 100 μm. **(B)** Absorption spectrum of recombinant Opn5m. Red line indicates fitted spectra. **(C)** Representative image series showing fluorescence changes during the excitation light exposure (450–490 nm) in melanotrophs of *opn5m*^**+/-**^ or *opn5m*^**-/-**^ medaka. Scale bar, 50 μm. **(D)** Time course of GCaMP fluorescence changes of melanotrophs in *opn5m*^**+/-**^ or *opn5m*^**-/-**^ medaka during exposure to the blue excitation light. *n* = 5 medaka. **(E), (F)** Response to the excitation light was suppressed in *opn5m*^**-/-**^ medaka compared to *opn5m*^**+/-**^ medaka, whereas response to CRH occurs similarly in *opn5m*^**+/-**^ and *opn5m*^**-/-**^ medaka. *n* = 5 medaka; *P* = 0.012, *P* = 0.31; Student’s *t* test. **(G)** Representative image series showing fluorescence changes during the excitation light exposure (450–490 nm) in HEK293A cells expressing mock or *opn5m*, and *gcamp6s*. Note that the cells were incubated with all-*trans*-retinal after transfection. Arrows indicate the cells transfected with *opn5m* and *gcamp6s*. Scale bar, 25 μm. **(H)** The maximum relative fluorescence of GCaMP during the whole recording (17.85 s) of each HEK293A cell expressing GCaMP6s (mock, 5cells) or cells expressing Opn5m and GCaMP6s (21cells). ***, *P* = 0.0005; Student’s *t* test. Data are represented as mean ± SEM.

Additionally, we elucidated the importance of direct photoreception by the pituitary even though the fish can acquire light information from the retina. We compared the [Ca^2+^]_i_ response of melanotrophs when white LED light was irradiated to the retina or the pituitary, using a semi-intact preparation with the whole brain, pituitary, and eye covered by the skull except for the ventral surface of the pituitary (Fig. 3A). This Ca^2+^ imaging indicated that light irradiation to the pituitary but not the retina induces a [Ca^2+^]_i_ increase in melanotrophs (Fig. 3B to D). Furthermore, to examine if natural sunlight can induce this [Ca^2+^]_i_ increase, we analyzed how much external light reaches the pituitary through the skull and the brain. From the analysis of light transparency using a semi-intact preparation retaining all structure above the pituitary (e.g., brain and skull roof), we estimated that approximately 2 to 4% external light reaches the pituitary in the wild strain of medaka (fig. S13A). Based on this transparency and the absorption spectrum of Opn5m (fig. S8), the white LED light used in this experiment at the intensity of 255 μmol m^-2^ s^-1^ is equivalent to the sunlight (∼2667 μmol m^-2^ s^-1^, measured on a sunny day) that reaches the pituitary in terms of activation of Opn5m (fig. S13B). Therefore, it can be concluded that the significant [Ca^2+^]_i_ increase observed in melanotrophs should be induced in their natural states. Furthermore, the transparency of the skull and the brain should be higher in larval stages. We performed Ca^2+^ imaging in larval (5–8 days post hatch) medaka using *ex vivo* preparation with intact whole head and found that this light response of melanotrophs also occurred in early stages (Fig. 3E and F). This demonstration in larval medaka suggests that this mechanism can be applied to other larger fishes in their larval stages.

**Fig. 3.**
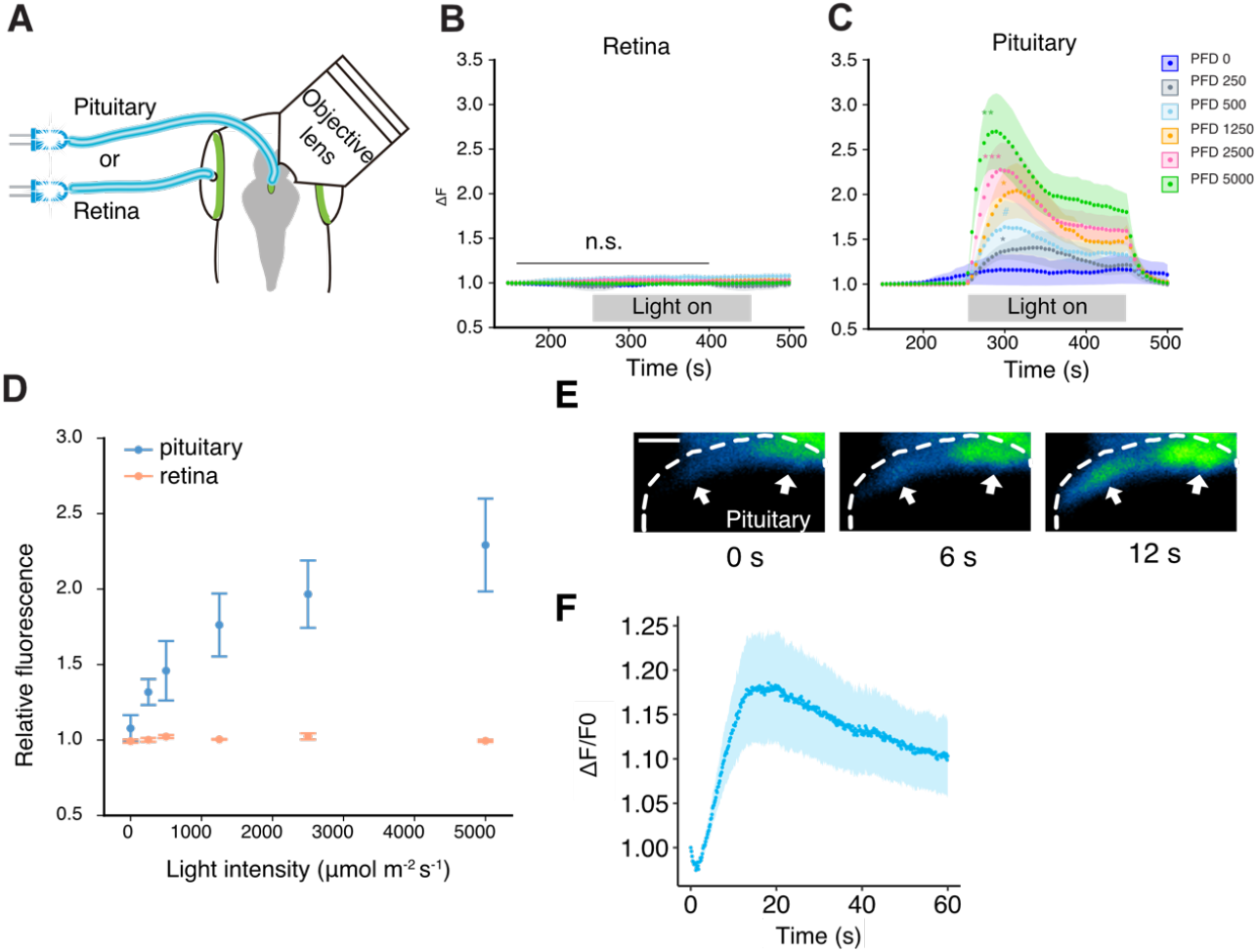
Direct photoreception in the pituitary is exclusively important for the Ca^2+^ response of melanotrophs. **(A)** The experimental scheme of Ca^2+^ imaging with local light irradiation using an optical fiber in semi-intact preparation. The retina or pituitary was separately irradiated with white LED. **(B-C)** Time course of GCaMP fluorescence changes of melanotrophs with local light irradiation to (B) retina or (C) pituitary. *n* = 7 medaka; ***, *P* = 0.0049; **, *P* = 0.006; *, *P* = 0.01, *P* = 0.01, #, *P* = 0.065, respectively; n.s., not significant; paired *t* tests were applied before and during irradiation. **(D)** Effect of LED irradiation of various light intensity to pituitary (blue) or retina (orange). Fold changes of GCaMP fluorescence of melanotroph are shown (during irradiation/before irradiation). **(E)** Representative image series showing GCaMP fluorescence changes of melanotrophs during the excitation light exposure (450–490 nm) in larva. The arrows indicate the melanotrophs. Note that fluorescence was observed from the intact head, through the skull and the brain. Fluorescence outside of the pituitary is autofluorescence. Scale bar, 12.5 μm. **(F)** Time course of GCaMP fluorescence changes of melanotrophs. *n* = 6 medaka. Data are represented as mean ± SEM.

To examine if this light-induced [Ca^2+^]_i_ increase in melanotrophs actually results in hormone release, we analyzed the contents of MSH and its derivatives in the medium where isolated pituitaries were incubated. During the incubation, 409 nm light irradiation or CRH was applied for 3 hours, and each medium was analyzed by liquid chromatography-mass spectrometry analysis (LC-MS). In the pituitary of WT medaka, but not in *opn5m*^-/-^ medaka, we observed an increase in the release of desacetyl α-MSH peptide by light stimulation. As expected, in both genotypes, CRH application increased desacetyl α-MSH as this effect is independent of Opn5m (Fig. 4A). Also, α-MSH showed an increasing trend by light stimulation in WT medaka, but not in *opn5m*^-/-^ medaka (fig. S14). Thus, as suggested by Ca^2+^ imaging experiments, light induces the release of hormones from melanotrophs via Opn5m. Finally, we tried to surmise the biological significance of this UV-induced release of MSH at the whole body level. Since it has been suggested that MSH increases the expression of *tyrosinase*, which is the rate-limiting enzyme in the melanogenesis pathway that is activated by MSH via Mc1r (*24, 25*), we analyzed the expression level of *tyrosinase* in *opn5m* KO medaka. Here, we used medaka under mixed LED light including 365 nm and normal white LED for at least 3 days. Based on the absorption spectrum of Opn5m, the spectra of sunlight, and this artificial light setup, this artificial light was estimated to have less than one-quarter the effect on Opn5m activation compared to the natural sunlight (fig. S15A). The *tyrosinase* expression level in the skin and the skull roof of *opn5m*^-/-^ medaka showed a suppressed *tyrosinase* expression level when compared to *opn5m*^+/-^ (Fig. 4B). In addition, the expression levels of both tyrosinase-related protein 1a (*tyrp1a*) and 1b (*tryp1b*), which are also involved in the melanogenesis pathway, were suppressed in *opn5m*^-/-^ medaka (fig. S15B). This evidence supports the notion that MSH, which is responsible for melanogenesis, is released to the general circulation via the activation of Opn5m. Taken together, this mechanism may play a role in the protection from harmful UV light that reaches as deep as the central nervous system.

**Fig. 4.**
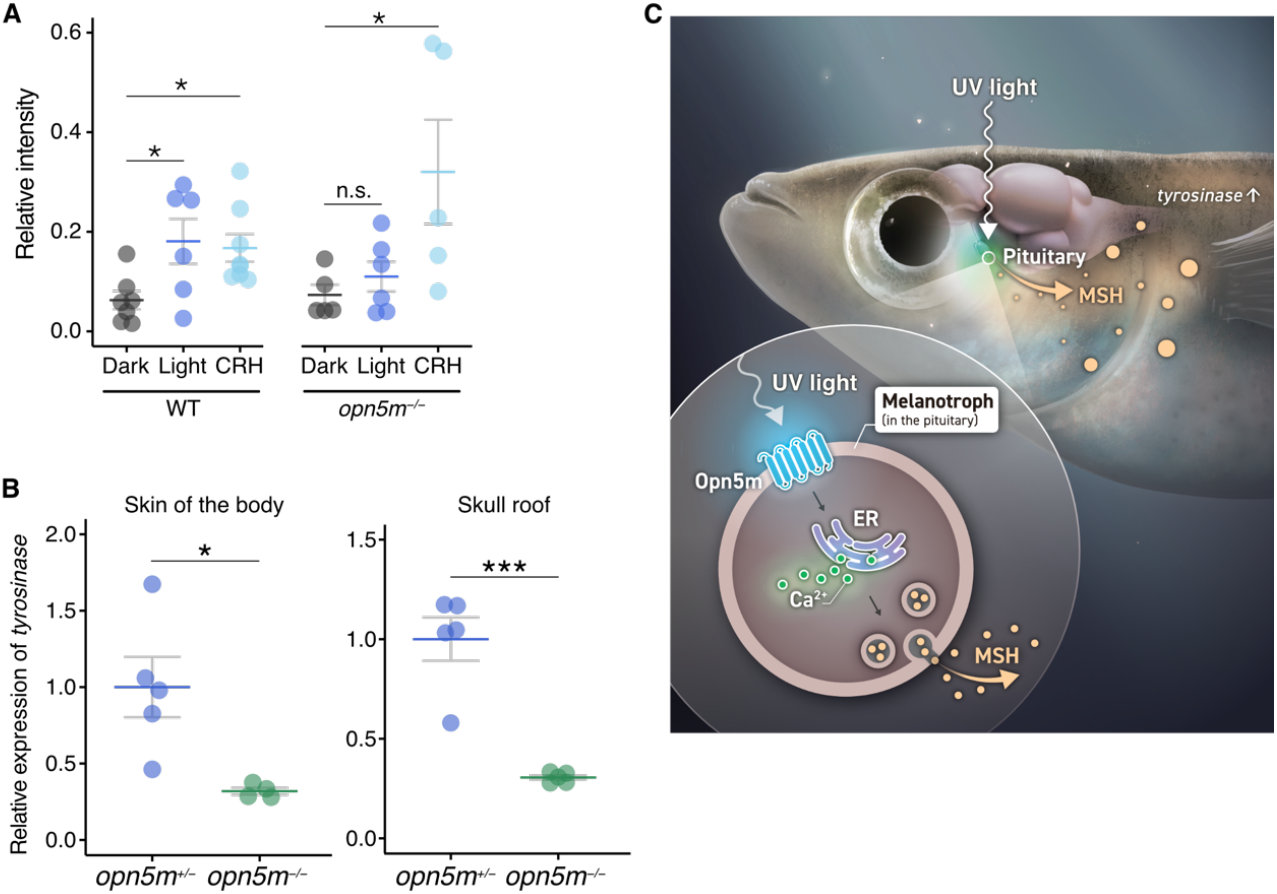
MSH is released by direct photoreception of melanotrophs, resulting in increased *tyrosinase* expression in the skin. **(A)** Effects of the light stimulation or CRH were analyzed in an *in vitro* preparation of isolated pituitaries. LC-MS analysis of the culture supernatant indicated that light induces desacetyl α-MSH peptide release in WT pituitaries, not but in *opn5m*^-/-^ pituitaries. Both WT and *opn5m*^-/-^ pituitaries show the CRH-induced MSH release. *n* = 7, 6, 6 and 5, 6, 5; *, *P* = 0.031, 0.041, 0.023; n.s., not significant; Dunnet tests. **(B)** The expression level of *tyrosinase* of *opn5*m^-/-^ medaka skin is lower than that of *opn5m*^+/-^ medaka. *N* = 5, 4 and *n* = 5, 5 medaka; ***,*P* = 0.00023; *,*P* = 0.019; Student’s *t* test. **(C)** Schematic illustration of the novel pathway of short-wavelength light induced MSH release. Sunlight, including short-wavelength light, reaches the pituitary melanotrophs. Opn5m, a non-visual photoreceptor expressed in the melanotrophs, receives UV light and induces Ca^2+^ release from endoplasmic reticulum. Increased intracellular Ca^2+^ triggers exocytosis in melanotrophs, which results in MSH release to the general circulation. Released MSH increases melanogenesis in the skin by increasing the expression of the rate-limiting enzyme, *tyrosinase*. Data are represented as mean ± SEM.

## Discussion

The present study identified a novel mechanism of the pituitary in which melanotrophs autonomously release hormones in response to short-wavelength light via Opn5m. This mechanism may contribute to the protection from invasive UV by enhancing melanogenesis (Fig. 4C). Among recent studies demonstrating the importance of non-visual opsin in various brain regions (*23, 26-28*), this is the first to note its existence in the pituitary, even deeper than the brain. In particular, the discovery that an endocrine cell of the pituitary directly perceives information outside of the body is surprising because the secretion of anterior pituitary hormones has been considered to primarily be regulated by the hypothalamus (*29, 30*). This finding significantly amplifies the potential roles of non-visual opsins, expanding their possible roles to a much wider range of organs than previously considered.

The discovery of the non-visual opsin involved in hormonal release to the general circulation in the present study also provides possibilities for future applications. For example, it could be a new optogenetic tool that enables the massive release of hormones or neurotransmitters. Recently, some non-visual opsins have demonstrated their possibility as tools for optogenetics in cultured cell lines and neurons in transgenic mice (*22*). Moreover, since the Opn5m localized in the pituitary is sensitive enough to be activated by transmitted sunlight and to increase [Ca^2+^]_i_, this sensitive opsin could theoretically be used to control the neurotransmitter release in any brain neurons theoretically. This suggests possible future application as a noninvasive optogenetic tool for free-moving small fishes, which may provide a paradigm shift in neuroscientific analyses at the individual level.

## Acknowledgments

We thank Mr. Satoshi Sekiguchi (University of Tokyo, Japan) for help with *pomc*:GCaMP construction. We are grateful to Drs. Susumu Hyodo, Shinji Nagata (University of Tokyo), Kanta Mizusawa (Kitasato University, Japan), and Yasuhisa Akazome (St. Marianna University, Japan) for their helpful discussion and comments. We also thank Drs. Yoshitaka Oka (University of Tokyo) and Kazuhiko Yamaguchi (RIKEN, Japan) for providing instruments. The authors thank Dr. Pamela Mellon (University of California–San Diego, La Jolla, CA, USA) for providing LβT2 cell line. We are grateful to Vesper Studio (Tokyo, Japan) for preparation of the schematic illustrations.

## Author contributions

Conceptualization: AF, KS, SK

Methodology: AF, KS, CF, CU, SK, AT, HO

Investigation: AF, KS, TY

Visualization: AF, KS, SK

Funding acquisition: AF, KS, SK

Project administration: SK

Supervision: SK

Writing – original draft: AF, KS, SK

Writing – review & editing: AF, KS, TY, CF, CU, SK

## Data and materials availability

All data are available in the main text or the supplementary materials.

## Supplementary Materials

Materials and Methods

Figs. S1 to S15

Tables S1 to S2

References (31–43)

## References and Notes

1. M. Andrabi, B. Upton, R. A. Lang, S. Vemaraju, An expanding role for nonvisual opsins in extraocular light sensing physiology. Annual Review of Vision Science 9, 1 (2023).

2. A. Terakita, The opsins. Genome Biology 6, 213 (2005).

3. B. A. Gheban, I. A. Rosca, M. Crisan, The morphological and functional characteristics of the pineal gland. Med Pharm Rep 92, 226–234 (2019).

4. S. Panda et al., Melanopsin (Opn4) Requirement for normal light-induced circadian phase shifting. Science 298, 2213–2216 (2002).

5. J. Falcón et al., Structural and functional evolution of the pineal melatonin system in vertebrates. Annals of the New York Academy of Sciences 1163, 101–111 (2009).

6. T. Ishikawa et al., in Laboratory Fish in Biomedical Research, L. d’Angelo, P. d. Girolamo, Eds. (Elsevier, Amsterdam, 2021), chap. 10, pp. 185-213.

7. C. Umatani, M. Nakajo, D. Kayo, Y. Oka, S. Kanda, in Laboratory Fish in Biomedical Research, L. d’Angelo, P. d. Girolamo, Eds. (Elsevier, Amsterdam, 2021), chap. 10, pp. 215-243.

8. S. Kanda, in Zebrafish, Medaka, and Other Small Fishes - New Model Animals in Biology, Medicine, and Beyond, H. Hirata, A. IIda, Eds. (Springer Science, 2018), pp. 99–111.

9. K. Murata, M. Kinoshita, Y. Kamei, M. Tanaka, K. Naruse, Medaka biology, management, and experimental protocols volume 2 preface. Medaka: Biology, Management, and Experimental Protocols, Vol 2 (2020).

10. T.-W. Chen et al., Ultrasensitive fluorescent proteins for imaging neuronal activity. Nature 499, 295–300 (2013).

11. D. O. Norris, J. A. Carr, in Vertebrate Endocrinology (Sixth Edition), D. O. Norris, J. A. Carr, Eds. (Academic Press, San Diego, 2021), pp. 151–204.

12. A. J. Bean, X. Zhang, T. Hökfelt, Peptide secretion: what do we know? The FASEB Journal 8, 630–638 (1994).

13. S. S. Stojilkovic, H. Zemkova, F. Van Goor, Biophysical basis of pituitary cell type-specific Ca<sup>2+</sup> signaling–secretion coupling. Trends in Endocrinology & Metabolism 16, 152–159 (2005).

14. S. Martens, H. T. McMahon, Mechanisms of membrane fusion: disparate players and common principles. Nature Reviews Molecular Cell Biology 9, 543–556 (2008).

15. K. Sato et al., Two UV-Sensitive photoreceptor proteins, Opn5m and Opn5m2 in ray-finned fish with distinct molecular properties and broad distribution in the retina and brain. PLoS One 11, e0155339 (2016).

16. T. Yamashita et al., Evolution of mammalian Opn5 as a specialized UV-absorbing pigment by a single amino acid mutation. J Biol Chem 289, 3991–4000 (2014).

17. T. Yamashita et al., Opn5 is a UV-sensitive bistable pigment that couples with Gi subtype of G protein. Proc Natl Acad Sci U S A 107, 22084–22089 (2010).

18. M. Jinek et al., A Programmable Dual-RNA guided DNA endonuclease in adaptive bacterial immunity. Science 337, 816–821 (2012).

19. T. N. Tran, J. N. Fryer, K. Lederis, H. Vaudry, CRF, Urotensin I, and sauvagine stimulate the release of POMC-derived peptides from goldfish neurointermediate lobe cells. General and Comparative Endocrinology 78, 351–360 (1990).

20. P. Thomas, P. L. Mellon, J. Turgeon, D. W. Waring, The L beta T2 clonal gonadotrope: a model for single cell studies of endocrine cell secretion. Endocrinology 137, 2979–2989 (1996).

21. K. Sato, T. Yamashita, H. Ohuchi, Mammalian type Opsin 5 preferentially activates G14 in Gq-type G proteins triggering intracellular calcium response. Journal of Biological Chemistry, 105020 (2023).

22. A. Wagdi et al., Selective optogenetic control of Gq signaling using human Neuropsin. Nature Communications 13, 1765 (2022).

23. K. X. Zhang et al., Violet-light suppression of thermogenesis by opsin 5 hypothalamic neurons. Nature 585, 420–425 (2020).

24. T. H. Nasti, L. Timares, MC1R, Eumelanin and pheomelanin: Their role in determining the susceptibility to skin cancer. Photochemistry and Photobiology 91, 188–200 (2015).

25. L.-m. Wang et al., The role of melanocortin 1 receptor on melanogenesis pathway in skin color differentiation of red tilapia. Aquaculture Reports 22, 100946 (2022).

26. K. Sato, K. N. Nwe, H. Ohuchi, The Opsin 3/Teleost multiple tissue opsin system: mRNA localization in the retina and brain of medaka (Oryzias latipes). Journal of Comparative Neurology 529, 2484–2516 (2021).

27. Y. Nakane et al., The saccus vasculosus of fish is a sensor of seasonal changes in day length. Nat Commun 4, 2108 (2013).

28. N. Nakao et al., Thyrotrophin in the pars tuberalis triggers photoperiodic response. Nature 452, 317–322 (2008).

29. G. Raisman, An urge to explain the incomprehensible: Geoffrey Harris and the discovery of the neural control of the pituitary gland. Annual Review of Neuroscience 20, 533–566 (1997).

30. G. Raisman, 60 years of neuroendocrinology: memoir: Geoffrey Harris and my brush with his unit. Journal of Endocrinology 226, T1–T11 (2015).

