## Supplementary Materials for "Direct photoreception of a pituitary endocrine cell, melanotroph, induces a hormone release"

**Materials and Methods:**

**Animals**

In the present study, we used medaka (*Oryzias latipes*). We used mature males as representatives unless otherwise mentioned (body weight: 0.07–0.24 g). All medaka were maintained under a 14-hour light and 10-hour dark photoperiod (light on at 0900–2300 or 0800–2200) at a water temperature of 27°C. These medaka were fed at least twice a day with live brine shrimp and flake food. All animals were maintained and used in accordance with the guidelines of the University of Tokyo for the use and care of experimental animals and the protocols have been approved by the Animal Care and Use Committee of the University of Tokyo (permission number P19-3, P22-7).

Generation of genetically modified medaka

Transgenic medaka

*pomc*:GCaMP transgenic medaka was generated as follows. As a region possibly harboring *pomc* enhancer and promoter, we used 3.7kb of 5' flanking region of POMC gene. This region was amplified from medaka genome with the primers (*pomc* UP3.7k_SE (backbonelink) and *pomc* UP_AS (GCaMPlink); table S1) and was fused with coding sequence of GCaMP6s (*10*). As described previously (*31*), larval stage-specific globin enhancer (*32*) fused with DsRed-Express was integrated to the construct for efficient screening of this transgenic medaka (fig. S1A). This construct was introduced to himedaka strain medaka by microinjection and the individual that has transgene was screened based on the fluorescence as described previously (*31*).

Generation of Knockout medaka

For the generation of knockouts for *opn5m* we used CRISPR/Cas9 system. Mixture of Cas9 Protein, tracr RNA, CRISPR RNA, GFP mRNA diluted with PBS and 0.02% phenol red (final concentration: Cas9 protein, 2 μg/μL; tracr RNA, 200 ng/μL; CRISPR RNA, 100 ng/μL; EGFP mRNA, 8 ng/μL) was injected into the embryos of *pomc*:GCaMP transgenic medaka (himedaka strain) or wild type medaka (d-rR strain). Two target sequences of guide RNA (gRNA) design including PAM (underlined) are as follows; CTTTGGGACTTCGTGCACGCTGG, TTTGATGCGGGCGTCGAAACTGG. CRISPR RNA and tracr RNA were synthesized by a commercial company (Integrated DNA Technology, Coralville, IA or Fasmac, Atsugi, Japan). After injection, GFP positive embryos were selected and then incrossed. F1 fish were then genotyped by using PCR amplification followed by sequence analyses with primers described in the primer list (table S1). Individuals that showed mutation in the target gene were crossed with fish with mutation in the same gene or wild type fish to produce homozygous/heterozygous transgenic offspring.

Specificity of GCaMP expression in *pomc* producing cells

In order to confirm the specificity of GCaMP expression in *pomc* producing cells, we performed dual labeling of *pomc* mRNA *in situ* hybridization (ISH) and GCaMP immunohistochemistry (IHC) on frozen section of the pituitary with the brain in *pomc*:GCaMP medaka. The medaka was deeply anesthetized with MS-222 (Sigma-Aldrich, Darmstadt, Germany) and quickly fixed by perfusion with 4% paraformaldehyde in PBS. Then dissected brain including pituitary was post-fixed with PFA for 1 hour and immersed in 30% (w/v) sucrose in PBS overnight. The brain-pituitary preparation was sagittally cryosectioned at 20 μm using a cryostat (CM3050S, Leica Microsystems, Welzlar, Germany) and mounted onto MAS-GP type A coated glass slides (Matsunami, Kishiwada, Japan). The GCaMP was labeled by anti- green fluorescent protein (EGFP) polyclonal antibody (#598, MBL, Nagoya, Japan) (diluted 1:1000 with PBS) for 4 hours. After labeled by biotinylated anti-rabbit IgG (Vector Laboratories, Burlingame, CA) for 1 hour, they were fixed with PFA for 5 minutes. These sections were then subjected to *in situ* hybridization protocol to visualize *pomc* mRNA expression with a standard protocol described in a previous study (*33*) using a digoxigenin (DIG)-labeled *pomc* probe used in a previous study (*34*). After ISH probe hybridization, we incubated the sections with anti-DIG POD (0.15 U/mL, Roche Diagnostics, Basel, Switzerland) for 1 hour and visualized *pomc* mRNA-expressing cells using TSA/Cy3 (diluted 1:100, PerkinElmer, Waltham, MA). After the *pomc* mRNA was detected, sections were incubated with ABC reagents (Vector Laboratories) for 1 hour and then Alexa Fluor 488 conjugated streptavidin for 1 hour (1:500 with PBS, ThermoFisher Scientific, Waltham, MA). Then sections were coverslipped with CC/Mount (Diagnostic BioSystems, Pleasanton, CA). The fluorescence was observed under a confocal laser-scanning microscope (FV-1000, Olympus, Tokyo, Japan).

**$Ca^{2+}$ imaging**

Intracellular $Ca^{2+}$ imaging with GCaMP

The fish were anesthetized by immersion in 0.02% tricaine methanesulfonate (MS-222, Sigma-Aldrich) and then decapitated. For *in vitro* $Ca^{2+}$ imaging of adults, the pituitary was thoroughly dissected out and was placed in a chamber with a fish artificial cerebrospinal fluid (ACSF) containing (in mM): NaCl 134, KCl 2.9, $CaCl_2$ 2.1, $MgCl_2$ 1.2, HEPES 10, and glucose 15 (adjusted to pH 7.4 with NaOH). For $Ca^{2+}$ imaging of larval medaka, we used *ex vivo* whole head preparation of 5-8 days post hatching was anesthetized with 0.02% MS-222, decapitated and the body was removed. Larva with their sex unidentified were used in this experiment. External recording solution was water containing the same anesthetic. GCaMP imaging was performed with an upright fluorescence microscope (Eclipse E600FN; Nikon, Tokyo, Japan) equipped with filter (EX; 450–490, Dichroic Mirror; 505, BA; 520 (LP)), and a light source, X-Cite 110LED Illumination System (Excelitas Technologies, Waltham, MA). During imaging experiments, the fluorescence images were captured (exposure: 50 ms; interval: 100-5,000 ms) by an electron multiplying-charge coupled device (EM-CCD) camera (QuantEM 512SC, Photometrics, Tucson, AZ) or a scientific complementary metal oxide semiconductor (sCMOS) camera (Andor Zyla 4.2 PLUS, Oxford Instruments, Belfast, UK). Acquisition of images were controlled by an imaging software, the Micro-Manager 1.4 (*35*). For quantification of the fluorescence, we analyzed 10 cells /individuals in adult medaka and 3 areas /individuals in larval medaka unless otherwise mentioned.

Examination of wavelength sensitivity of the melanotrope

To examine the effects of stimulation lights of different wavelength and intensity, we illuminated stimulation light only in the interval of imaging to avoid that the stimulation light is directly sensed by camera. For the precise control of this, we used a single-board microcontroller, Arduino UNO R3 (Arduino, Turin, Italy) with a simple script and a solid-state relay, G3TB-ODX03P DC5-24 (Omron, Kyoto, Japan) (fig. S3). Since continuous robust $Ca^{2+}$ rise in the melanotropes did not occur during the recording with more than 5 second interval (fig. S4A), we recorded the $Ca^{2+}$ imaging with 5-second cycles, where each cycle involved fluorescence image capture of 50 ms, followed by an interval of approximately 4950 ms. Light irradiations as stimulation light for 4800 ms were applied during this recording interval (fig. S4B). Irradiation protocol of stimulation light started 200s after recording without stimulation light because transient increase of $Ca^{2+}$ by the excitation light for imaging occurred within the first 200 s and later it gets steady state that allows us to evaluate the effect of additional stimulation.

The stimulation lights were irradiated through the optical fiber (plastic fibers for normal experiments or quartz fibers for experiments including UV light). We prepared the pituitary preparation as described above and located the fiber in the close vicinity of the pituitary. The LED light we used was 365 nm (LED generic, Minami-alps, Japan), 420 nm (OSG5XNE1C1E), 450 nm (OSB5XNE1C1E), 520 nm (OSY5XNE1C1E), 590 nm (OSG5XNE1C1E), 630 nm (OSR5XNE1C1E), 740 nm (LED generic). The light intensity was measured by the PG200N Spectral PAR Meter (UPRtek, Zhunan Township, Taiwan) in all subsequent experiments.

$Ca^{2+}$ responsiveness in melanotrope to the photoreception in the retina and pituitary

To compare the effect of light from retina and pituitary on melanotrope, we prepared a semi-intact preparation with whole brain, pituitary and eye covered by skull except for the ventral surface of the pituitary. During $Ca^{2+}$ imaging acquisition of pituitary melanotrope as described above, stimulation light of various strength through optical fiber was irradiated to the retina or pituitary. Note that the stimulation light was from a white color LED with broad spectrum (LED generic), and was illuminated during the 5 s intervals of $Ca^{2+}$ imaging acquisitions as described above.

$Ca^{2+}$ imaging of dissociated single melanotrope

To examine if a single melanotrope show sensitivity to light, we performed single cell $Ca^{2+}$ imaging using dissociated melanotrope of *pomc*:GCaMP medaka pituitary. First, we thoroughly dissected out the pituitary from 1-3 male or female *pomc*:GCaMP medaka as described above. The pituitaries were incubated with collagenase (150 U/mL) in 500 µL L-15 medium (Wako, Osaka, Japan) with 1× penicillin-streptomycin for 30 min 26°C. Then the posterior parts of the pituitaries were dissociated by pipetting with a pipette tip with its tip sharply processed (~100µm in diameter). Using these dissociated melanotropes labeled with GCaMP, we performed $Ca^{2+}$ imaging under the upright fluorescence microscope set up described above (exposure: 50ms, interval: 100ms). We performed 5 independent experiments, analyzing 4–9 GCaMP-expressing cells in each experiment.

Pharmacological analysis of intracellular $Ca^{2+}$ signaling

To identify the source of increasing $[Ca^{2+}]_i$, we used $Ca^{2+}$ free ACSF (in mM): NaCl 134, KCl 2.9, $MgCl_2$ 1.2, HEPES 10, and glucose 15 (adjusted to pH 7.4 with NaOH), or normal ACSF containing 100 µM $CdCl_2$ or 500 µM 2-aminoethoxydiphenyl borate (2-APB, Wako). We performed $Ca^{2+}$ imaging (exposure: 50 ms, interval: 100 ms) for 30 seconds to evaluate the $Ca^{2+}$ response to the excitation light just after applying each solution as follows: normal ACSF (before) for approximately 3 min, $Ca^{2+}$ free ACSF or normal ACSF containing one of the drugs (drug application) for 20 min, normal ACSF (washout) for 30 min. The normalized $[Ca^{2+}]_i$ response was calculated by dividing the maximum fluorescence during application by the intensity of the initial fluorescence. For perfusion of the solutions, a peristaltic pump (Rainin Dynamax RP-1, Rainin, Columbus,

OH) was used.

Ca$^{2+}$ responsiveness of melanotrophs to corticotropin releasing hormone (CRH) in *opn5m*$^{+/-}$ or *opn5m*$^{-/-}$

To examine if the response of *opn5m*$^{-/-}$ pituitary is normal other than that to light, we performed Ca$^{2+}$ imaging with CRH perfusion (CRF, Ovine, Peptide institute, Osaka, Japan). CRH peptide was diluted to 400 nM in ACSF. The fluorescence images of GCaMP were recorded (exposure: 50 ms; interval: 5 s) and CRH perfusion was stared at least 200 s after recording started, after GCaMP fluorescence of melanotrophs had become stable. After the CRH application, the pituitary was washed out with ACSF for 10 min. We analyzed the fluorescence intensity for each for statistical analysis: average intensity of 100 s before CRH perfusion (before), average intensity of the last 100 s of CRH perfusion (CRH), and average intensity of last 100 s of ACSF washout (wash). Relative GCaMP fluorescence was calculated by dividing the average intensity during CRH perfusion by before CRH application.

Examination of Opn5m activity in cultured cells

To examine the ability of Opn5m to increase $[Ca^{2+}]_i$ in response to the light, we utilized heterologous expression system using HEK293A cells and LβT2 cells. HEK293A cells and LβT2 cells were cultured in DMEM (Wako) with 10% heat-inactivated fetal bovine serum (FBS) and 1× penicillin-streptomycin in a $CO_2$ incubator 37°C. In ~70% confluence, cells were transferred to poly-L-lysine coated coverslips, and Opn5m expression vector (ENSORLT00000019589.2) or mock (2.7 μg) and GCaMP6s plasmid (0.9 μg) were transfected to HEK293A cells with PEI (23 μg) in Opti-MEM (360 μL) for overnight. In the case of transfection to LβT2 cells, Opn5m expression vector (0.3 μg) and GCaMP6s plasmid (0.06 μg) were transfected with 2 μL Lipofectamine 2000 (Invitrogen, Waltham, MA) in Opti-MEM (300 μL) for 4 hours. After that, we changed the medium containing all-*trans*-retinal or 11-*cis* retinal (final concentration, 5 μM) and further incubated for 1-24 hours under a dark condition. For the Ca$^{2+}$ imaging experiment, the coverslip was placed in the chamber filled with ACSF (exposure: 50 ms, interval: 100 ms). Because both all-*trans*-retinal and 11-*cis* retinal gave similar results, we mostly used all-*trans*-retinal due to its easy availability.

**Analysis of light transmittance rate to the pituitary through the skull**

To estimate the transmittance rate of various wavelengths of light to the pituitary through the skull and brain, we prepared a semi-intact preparation consist of skull, whole brain, and pituitary by removing all body parts ventral to the pituitary. Note that we used Kiyosu strain here as a wild strain of medaka. From a light source (U-HGLGPS, Olympus, Tokyo, Japan), light with specific wavelength was acquired using bandpass filters 370 nm, 400 nm, 450 nm, 500 nm, 550 nm, 600 nm, 650 nm, 700 nm, or 750 nm (FKB-VIS-10, Visible Bandpass Filter, Thorlabs, NJ) equipped in a filter wheel (#34-545; Edmund Optics, Cranbury, NJ). The light was introduced to the stereomicroscope (M165FC, Leica Microsystems) with optical fiber and transmitted images with or without the semi-intact preparation were taken (Zyla 4.2 Plus sCMOS; Micro-Manager 2.0-γ). From the captured images, the brightness of the pituitary area was measured by ImageJ. The transmittance rate was calculated by brightness of the picture area with preparation / brightness of the same area without preparation. Slight difference in the overall intensity was further calibrated by the intensity of background regions.

**Estimation of the strength of white LED that is equivalent to sunlight in terms of Opn5m activation**

The strength of white LED light equivalent to the sunlight in terms of Opn5m activation was calculated as follows.

$$\frac{(\text{intensity of total white LED}) * \int_{350}^{500} (\text{intensity of sunlight at } x \text{ nm}) * (\text{relative absorption of Opn5m spectrum at } x \text{ nm}) * (\text{transparency at } x \text{ nm})\, dx}{\int_{350}^{500} (\text{intensity of white LED at } x \text{ nm}) * (\text{relative absorption of Opn5m spectrum at } x \text{ nm})\, dx}$$

Spectrum of the light and absorption spectrum of Opn5m are shown in fig. S13A and Fig.2B, respectively. Transparency is shown in fig. S13B. As transparency was measured in several representative wavelengths, these representative values were applied to the neighboring wavelengths.

**Expression, purification, and spectrophotometry of recombinant medaka Opn5m**

The cDNA of medaka Opn5m was tagged by the epitope sequence of monoclonal antibody Rho1D4 (ETSQVAPA) at the C terminus and was inserted into the mammalian expression vector pCAGGS. The plasmid DNA was transfected into HEK293S cells using the calcium phosphate method. Culture medium was replaced with fresh one containing 5 μM 11-*cis*-retinal one day after transfection. After one more day kept in the dark, the cells were collected and solubilized with 1% n-dodecyl-β-d-maltoside (DDM) in buffer A (50 mM HEPES, pH 6.5, and 140 mM NaCl). The extracted medaka Opn5m was applied to Rho1D4-conjugated agarose. The medaka

Opn5m pigment was eluted with buffer A containing 0.02% DDM and the synthetic peptide of the Rho1D4 epitope sequence.

Absorption spectra were recorded at 0°C with a Shimadzu UV-2450 spectrophotometer. The sample was irradiated with UV light through a UV-D35 glass filter (Asahi Technoglass, Haibara, Japan), or with yellow light through a Y-49 cutoff filter (Toshiba, Tokyo, Japan) from a 1 kW projector lamp (Rikagaku seiki, Tokyo, Japan).

**Chromogenic *in situ* hybridization**

Detection of *opn5m*, *pomc*, *crhr1*, and *crhr2* with conventional chromogenic *in situ* hybridization was performed according to Sato *et al.*, 2021(*26*). The probes for *opn5m* were as described previously (*26*). To prepare the probe templates, the cDNAs of *pomc, crhr1* and *crhr2* (NCBI accession number; XM_004066456.2, GFIO01006283.1, GFIO01012195.1) were amplified using the primers, listed in table S1.

**Fluorescence *in situ* hybridization with SABER-FISH**

To detect mRNA of *opn5m* and *pomc* simultaneously, we used fluorescence *in situ* hybridization with signal amplification by exchanging reaction (SABER-FISH). Probe design for medaka *opn5m* and *pomc* was performed using OligoMiner (*36*) software with medaka genomic assembly ASM223467v1. Probe and branch oligo extensions by primer exchanging reaction were carried out according to the original paper for SABER-FISH (*37*) with the oligo DNAs listed in table S2. In the SABER-FISH, signal of *opn5m* was amplified by two iterations of branching reactions. Branching amplification was not applied to detection of *pomc*. The SABER-FISH was performed as follows with slight modification from the original protocol. Medaka brain samples were fixed overnight in 4% PFA/PBS at 4°C after dissection. After cryoprotection by immersion in 20% sucrose/PBS, the tissues were sectioned at 15 μm, adhered on the slide glass, and dried completely before use. The tissue sections on slide glass were rehydrated by PBS, treated with 0.05 mg/mL pepsin in 0.01 M HCl at 37°C for 20 min, rinsed twice with PBST. After incubation with Whyb (40% deionized formamide, 2xSSC, 1% Tween-20) at 43°C for 10 min, the probes were applied to the sections at 2 μg/mL in Hyb1 (40% deionized formamide, 10% Dextran sulfate (Sigma-Aldrich D8906), 2xSSC, 1% Tween-20) for 16 hours at 43°C. After hybridization of probes, the sections were rinsed with Whyb once and washed twice 30 min with Whyb, twice 5 min with 2xSSCT 43°C. The sections were rinsed with PBST twice at RT. For signal amplification with branching oligo, branches were applied to the sections at 2 μg/mL in

Hyb1 for 6-12 hours at 37°C. Wash was performed similarly as done after probe hybridization. After two rounds branch hybridizations for detection of *opn5m*, fluorophore conjugated oligos were applied at 0.2 μM in PBST for one hour at 37°C. Then, the sections were washed twice 5 min with PBST at 37°C, counterstained with Hoechst33342 (1 μg/mL), coverslipped with homemade polyvinyl alcohol/glycerol mounting medium, and imaged with a confocal laser-scanning microscope (LSM 780, Carl Zeiss Microscopy GmbH, Oberkochen, Germany).

**Hormone detection secreted from medaka pituitary by liquid chromatography-mass spectrometry**

Wild type and *opn5m*$^{-/-}$ adult female medaka (d-rR strain) were used. Sampling of pituitaries were started at approximately at 15:00 (ZT7). Except for three hours incubation with or without violet light and CRH, all the procedure were conducted under ambient light. Three medaka were anesthetized with 0.04% MS-222, decapitated, and pituitary glands were harvested and collected in ice-cold ACSF. Three pituitaries had been spined-down in a plastic tube. "Dark" sample tube was covered with tin foil and incubated at 28°C for 3 hours. "Light" sample tube was irradiated from the bottom of the tube by light plate apparatus equipped with violet LED (409 nm peak wavelength, 40 μmol m$^{-2}$ s$^{-1}$ photon flux density) for three hours at 28°C. The "CRH" sample was kept in the dark at 28°C for three hours in ACSF containing 0.4 μM ovine CRH. After three hours incubation at 28°C, the supernatants were collected, spiked with 1 ng [Nle$^4$,D-Phe$^7$]-α-MSH as an internal standard for relative quantification, and desalted by MonoSpin C18 (GL Sciences, Tokyo, Japan) according to the manufacturer's instruction. The desalted sample was separated on L-column ODS (0.1 id x 150 mm, 3 μm particle size; CELI) using a gradient of solvent A (97.9% $H_2O$, 2% MeCN, 0.1% HCOOH) and solvent B (9.9% $H_2O$, 90% MeCN, 0.1% HCOOH). The gradient program was 15% of solvent B in 0 min, 25% of solvent B in 15 min, and 90% of solvent B from 30 to 40 min at a flow rate of 0.5 μL min$^{-1}$. The effluent peptides from the HPLC were directly infused into an electrospray ion source of an LTQ-Orbitrap Discovery mass spectrometer (ThermoFisher Scientific). MS and MS$^2$ spectra were obtained using FTMS and ion trap analyzers, respectively. All of the ions were detected in positive mode. For the relative quantification, mass chromatograms of *m/z* 549.6194 ± 0.005 for $[M+3H]^{3+}$ of [Nle$^4$,D-Phe$^7$]-α-MSH, *m/z* 555.6049 ± 0.005 for $[M+3H]^{3+}$ of α-MSH, *m/z* 541.6014 ± 0.005 for $[M+3H]^{3+}$ of desacetyl α-MSH, and *m/z* 569.6084 ± 0.005 for $[M+3H]^{3+}$ of diacetyl α-MSH were excerpted. The relative abundance of the peptides were calculated by the division of the peak area intensity of α-MSH, desacetyl α-MSH, or diacetyl α-MSH by that of [Nle$^4$,D-Phe$^7$]-α-MSH in each sample. Those identities of [Nle$^4$,D-Phe$^7$]-α-MSH, α-MSH, desacetyl α-MSH, diacetyl α-MSH were confirmed by corresponding MS2 fragmentation spectra. LC-MS data were analyzed using MZmine3 (*38*) and IgorPro ver 9.02 software.

**Quantitative RT-PCR of *tyrosinase* in the skin**

Generated knockouts and their heterozygote siblings were kept under the light condition including UV light (UV LED (HCRave-TG0001, Gao chuang) and white LED (DS-UV365E-K1OC-M, Everfine)) for at least 3 days (14:10 L:D cycle). The light strength at each wavelength was lower than the sunlight (fig. S15A). Here, we used randomly mixed male and female medaka. Sampling of skin was started at approximately at 18:00 (ZT9). For real-time PCR analysis, these medaka were anesthetized, and their skin were peeled off from the body surface excluding the head. Also, the skull roofs were sampled. Each skin was homogenized in a tube by micro-smasher (TOMY SEIKO, Tokyo, Japan) with the lysis buffer for RNA extraction for 45 s. Total RNA was extracted from each skin using Fast Gene RNA Basic kit (Nippon Genetics, Tokyo, Japan) according to the manufacturer's protocol. Genomic DNA was removed by DNaseI (Invitrogen or Nippon Gene, Tokyo, Japan) treatment. Total RNA was reverse transcribed with PrimeScript RT kit (Takara, Kusatsu, Japan) according to the manufacturer's instructions. The cDNA was subjected to quantitative PCR (qPCR) using KAPA SYBR fast qPCR kit (Nippon Genetics) with LightCycler 480 II system (Roche Diagnostics). The temperature profile of the reaction was 95°C for 5 min, 45 cycles of denaturation at 95°C for 10 s, annealing at 60°C for 10 s, and extension at 72°C for 10 s. The PCR products was verified by melting curve analysis. The data was normalized by a housekeeping gene, β-actin (*actb*). The primer pairs used in the real-time PCR are shown in Supplemental table (table S1).

**Data analysis**

All values are shown as the mean ± SEM. For analysis and calculation of the fluorescence intensity changes in the melanotropes of $Ca^{2+}$ imaging, we used ImageJ (FIJI, National Institutes of Health, Bethesda, MD). Statistical analyses with two groups were examined by Student's *t* test, while groups of more than three were examined by Dunnett's test unless otherwise explained. Statistical analyses were performed with R Studio (version 1.2.5033) (*39*) or Kyplot5.0 (KyensLab, Tokyo, Japan).

Fig. S1

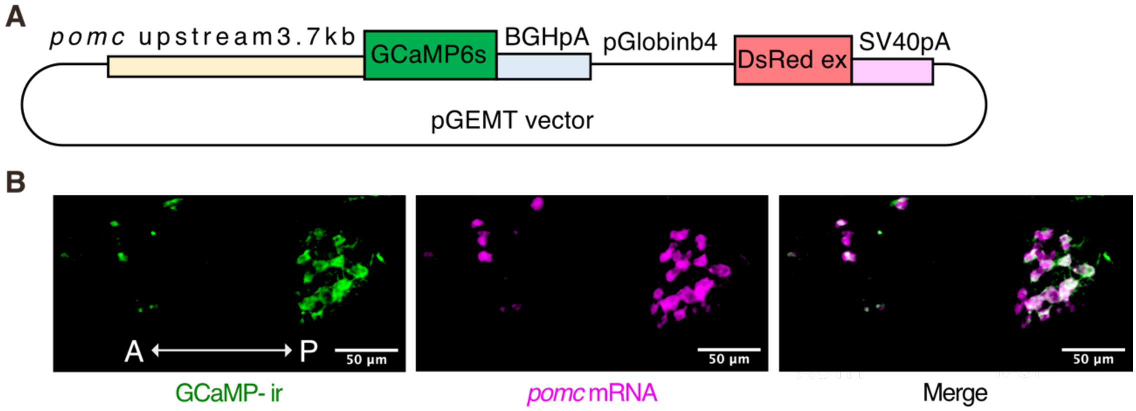


**Fig. S1. Generation of *pomc*:GCaMP6s transgenic medaka.**

**(A)** The construct used to generate *pomc*:GCaMP6s medaka. **(B)** Double labeling of EGFP (GCaMP) immunohistochemistry and *pomc in situ* hybridization. Based on previous studies (*11, 40*), corticotrophs (rostral pars distalis, anterior: A) and melanotrophs (pars intermedia, posterior: P) were identified by their localization in the pituitary. In corticotrophs, 98% of GCaMP-immunoreactive cell bodies were *pomc* mRNA–positive in the pituitary. In melanotrophs, 94% of GCaMP-immunoreactive cell bodies were *pomc* mRNA–positive in the pituitary. Scale bar, 50 µm.

Fig. S2

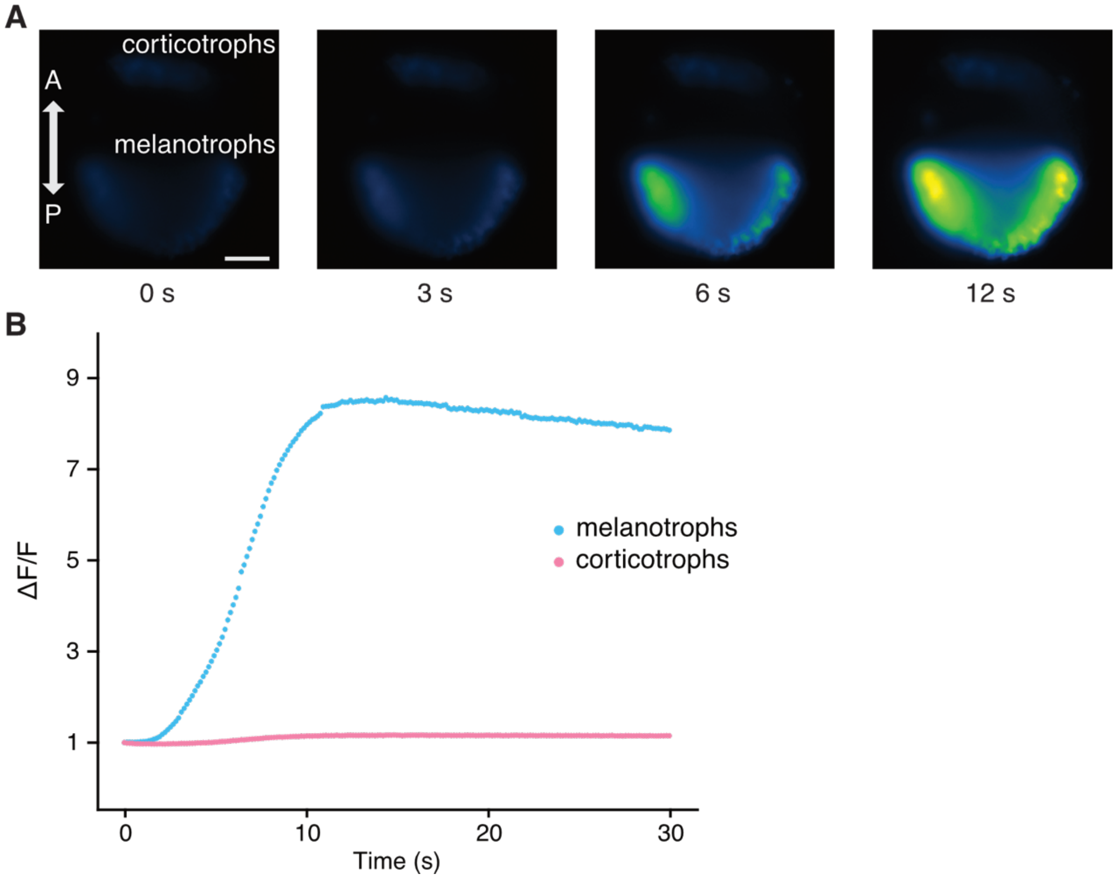


**Fig. S2. Melanotrophs in the pituitary show a [$Ca^{2+}$]$_i$ increase during the fluorescence observation under the blue excitation light in a whole brain pituitary preparation.**
**(A)** Representative image series of the pituitary of *pomc*:GCaMP medaka showing GCaMP fluorescence change during the fluorescence observation with excitation light exposure (450–490 nm). Note that the pituitary is connected to the brain in this preparation. Scale bar, 50 µm.
**(B)** Time course of GCaMP fluorescence change of melanotrophs and corticotrophs (average intensity of 10 cells).

**Fig. S3**

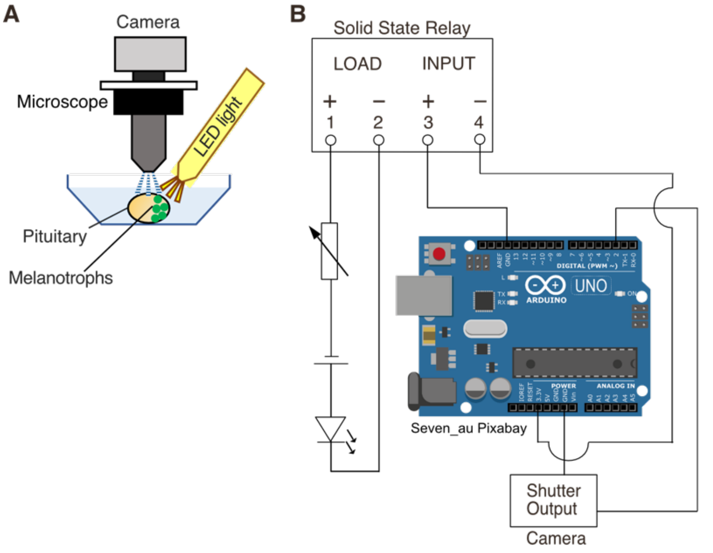


C

```
int waiting = 1;
int illumi = 0;
void setup(){
  Serial.begin(9600);
  pinMode(2, INPUT);
  pinMode(11, OUTPUT);
}
void loop() {
 int status =  PIND & _BV(2);
 Serial.println(status);
    if(status >3){
 Serial.println(illumi);
delay(50);
 if (illumi >= 40 && illumi < 52){
      PORTB  |= _BV(3) ;
 }
 else
 {
  PORTB &= ~_BV(3) ;
 }
 delay(4800);
       PORTB &= ~_BV(3) ;
       illumi++;
}
}
```

**Fig. S3. $Ca^{2+}$ imaging experimental setup for evaluation of the stimulation light.** **(A)** The experimental scheme of $Ca^{2+}$ imaging with LED stimulation during $Ca^{2+}$ imaging. **(B)** Irradiation timing of the stimulation light was controlled by a single-board microcontroller, Arduino, with a solid-state relay. The voltage of the DC power supply was altered by the built-in switch (3–12 V) and the variable resistor shown in the electrical diagram. When a single photo is taken, the camera sends a signal to Arduino. Arduino turns on the stimulation light after a delay (50 ms), then turn on after 4800 ms, which avoids the simultaneous occurrence of stimulation and image acquisition. **(C)** The script ran on Arduino that enables the mutual exclusion of fluorescence recording and stimulation light irradiation.

**Fig. S4**

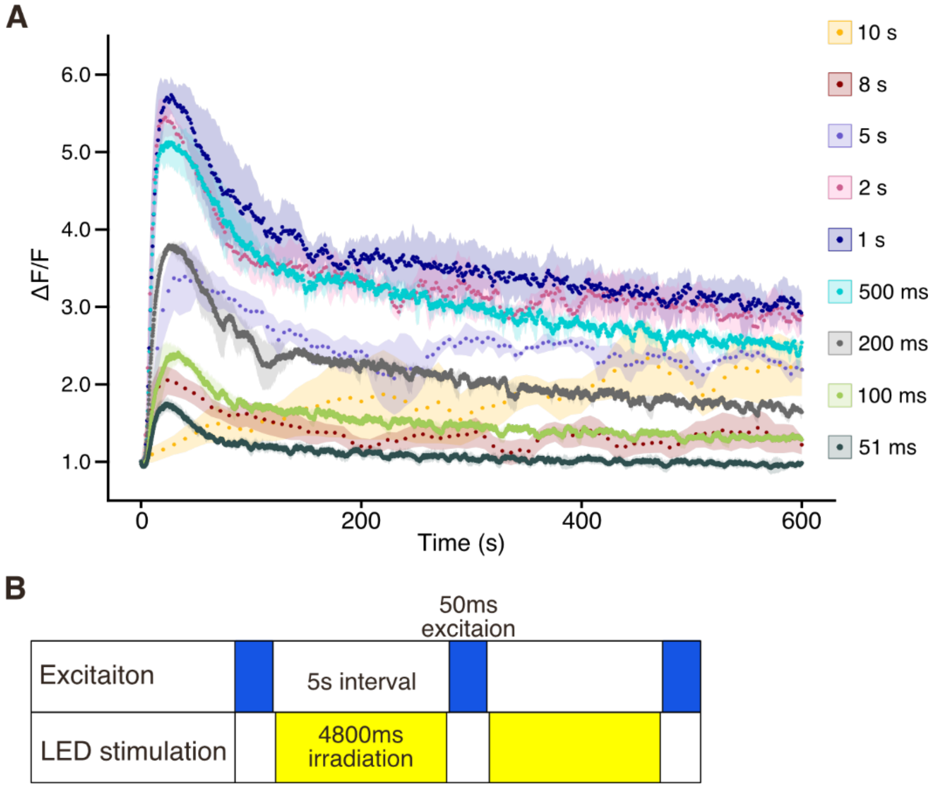


**Fig. S4. After continuous recordings with 5 s interval image acquisitions for 200 s, $[Ca^{2+}]_i$ becomes low and steady, which allows evaluation of the effect of additional stimulation .** **(A)** For the $Ca^{2+}$ imaging examining the effect of stimulation light in the presence of excitation light, it was necessary to find conditions with an appropriate interval time to avoid continuous elevation of $[Ca^{2+}]_i$ or breaching caused by the excitation light of $Ca^{2+}$ imaging. $Ca^{2+}$ imaging with 50 ms excitation was examined with various intervals of image acquisitions. The graph shows the GCaMP fluorescence changes in melanotrophs during $Ca^{2+}$ imaging with different intervals (51 ms, 100 ms, 200 ms, 500 ms, 1 s, 2 s, 5 s, 8 s, 10 s; $n = 3$). This suggests that after long continuous acquisition, the fluorescence level becomes lower and stable. In case of 5 s intervals, which is convenient for irradiation during the interval, it provides stable fluorescence level after 200 s recording. **(B)** Experimental scheme of GCaMP image acquisition and light stimulation controlled by Arduino script. Note that image acquisition and light stimulation is performed with mutual exclusion.

**Fig. S5**

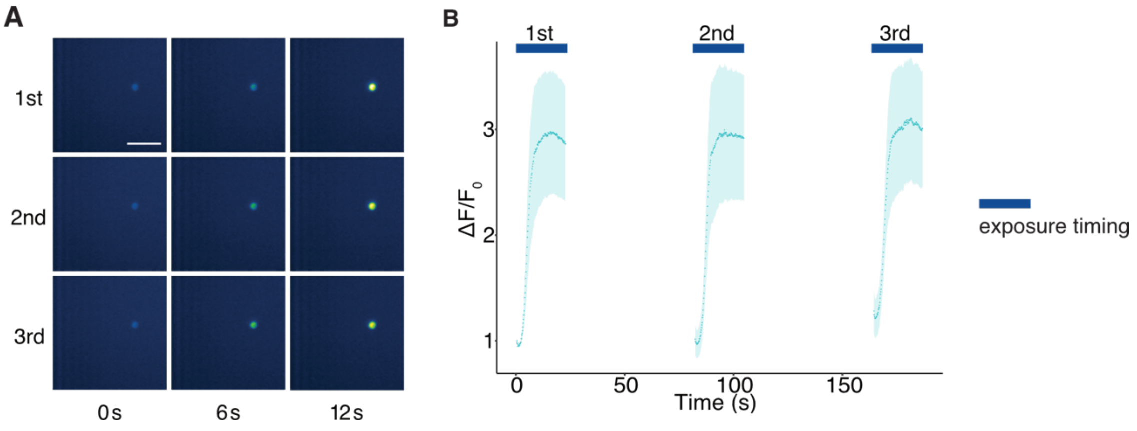


**Fig. S5. An isolated melanotroph responds to the excitation light repetitively.**

**(A)** Representative GCaMP fluorescence images of an isolated melanotroph in three repetitive $Ca^{2+}$ imaging trials. Each row represents each trial. In the $2^{nd}$ and $3^{rd}$ trials, the initiation of each trial is represented as 0 s. Scale bar, 50 µm. **(B)** Three repetitive trials similarly increased fluorescence intensity (60 s interval, 12 cells from 4 individuals).

**Fig. S6**

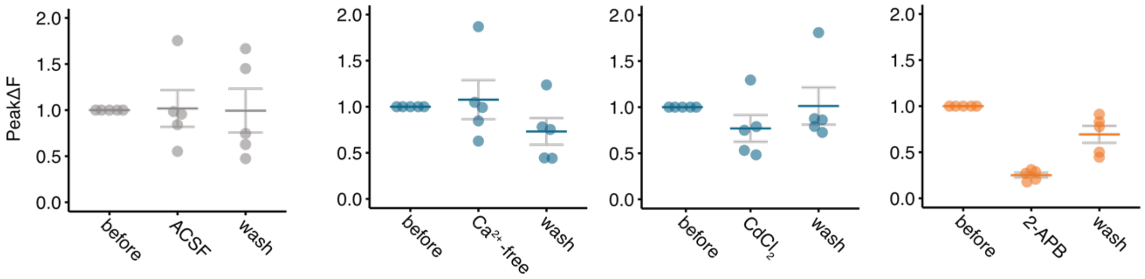


**Fig. S6. Light-induced $[Ca^{2+}]_i$ increase is derived from endoplasmic reticulum calcium store via IP3 receptor.**

The peak amplitudes of GCaMP fluorescence changes in melanotrophs before, during, and after washout are shown for each drug application. These data are the original data from Fig. 1F, which was calculated and statistically analyzed in the main text. These original data indicate that the effect of drug application was completely diminished after washout. *n* = 5 medaka.

**Fig. S7**

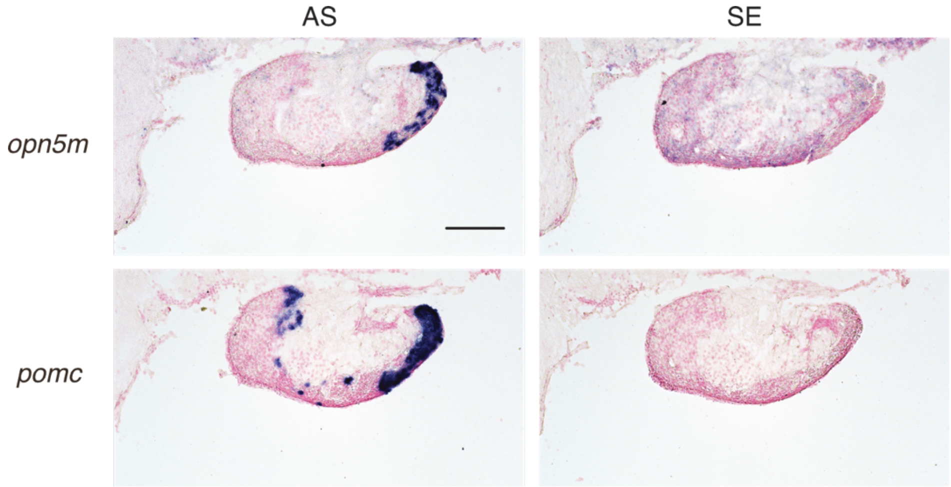


**Fig. S7. *opn5m* is expressed in the area where melanotrophs localize in the medaka pituitary.**

Representative photographs of *in situ* hybridization of *opn5m* in the pituitary. Only sections hybridized with anti-sense (AS) probe are labeled for both *opn5m* and *pomc*. These data suggest that a non-visual photoreceptor, *opn5m*, is expressed in the area where the caudal population of *pomc* expressing cells, melanotrophs, localize. Scale bar, 100 μm.

**Fig. S8**

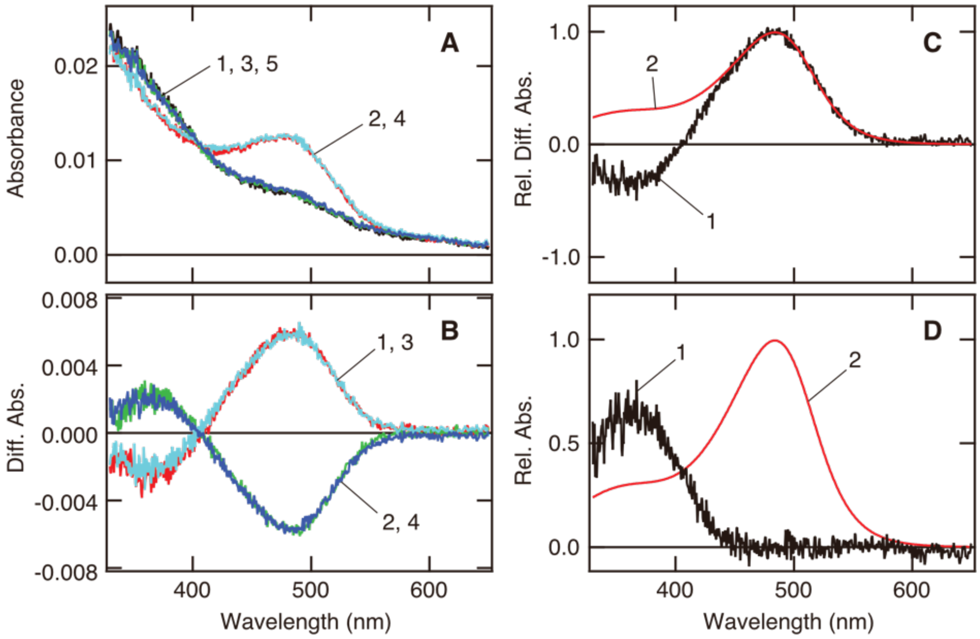


**Fig. S8. Medaka Opn5m absorption spectrum**

**(A)** Absorption spectra of purified medaka Opn5m bound to 11-*cis*-retinal before and after light irradiations. Spectra were recorded before irradiation (curve 1, black), after UV light (350 nm) irradiation (curve 2, red), after subsequent yellow light (> 470 nm) irradiation (curve 3, green), after UV light re-irradiation (curve 4, cyan) and after yellow light re-irradiation (curve 5, blue). **(B)** Difference spectra of medaka Opn5m calculated based on the absorption spectra in panel A. Curves 1to 4 in this panel show the difference spectrum calculated by subtracting curve 1 from 2, 2 from 3, 3 from 4 and 4 from 5 in panel A. **(C)** Calculation of the absorption spectrum of medaka Opn5m based on the difference and model spectra. Curve 4 in panel B was inverted and scaled to be 1.0 at the positive maximum (curve 1, black). The spectral region longer than the maximum of curve 1 was fitted with a model spectrum (curve 2, red) according to Lamb and Govardovskii *et al*. (*41, 42*). Subtraction of curve 1 from 2 leads to curve 1 in panel D. **(D)** The calculated absorption spectra of 11-*cis*- (curve 1, black) and all-*trans*-retinal-bound (curve 2, red) forms of medaka Opn5m. Curve 2 (red) is identical to curve 2 in panel C. The previous studies showed that 11-*cis*- and all-*trans* retinal bound forms are interconvertible by light irradiations (*16, 43*) Thus, curves 1 and 2 correspond to 11-*cis*- and all-*trans*-retinal bound forms, respectively. The absorption maxima of those 11-*cis*-retinal-bound and all-*trans*-retinal-bound forms are 363 and 484 nm. The curve 1 in panel D is identical to the black curve in Fig. 2B.

**Fig. S9**

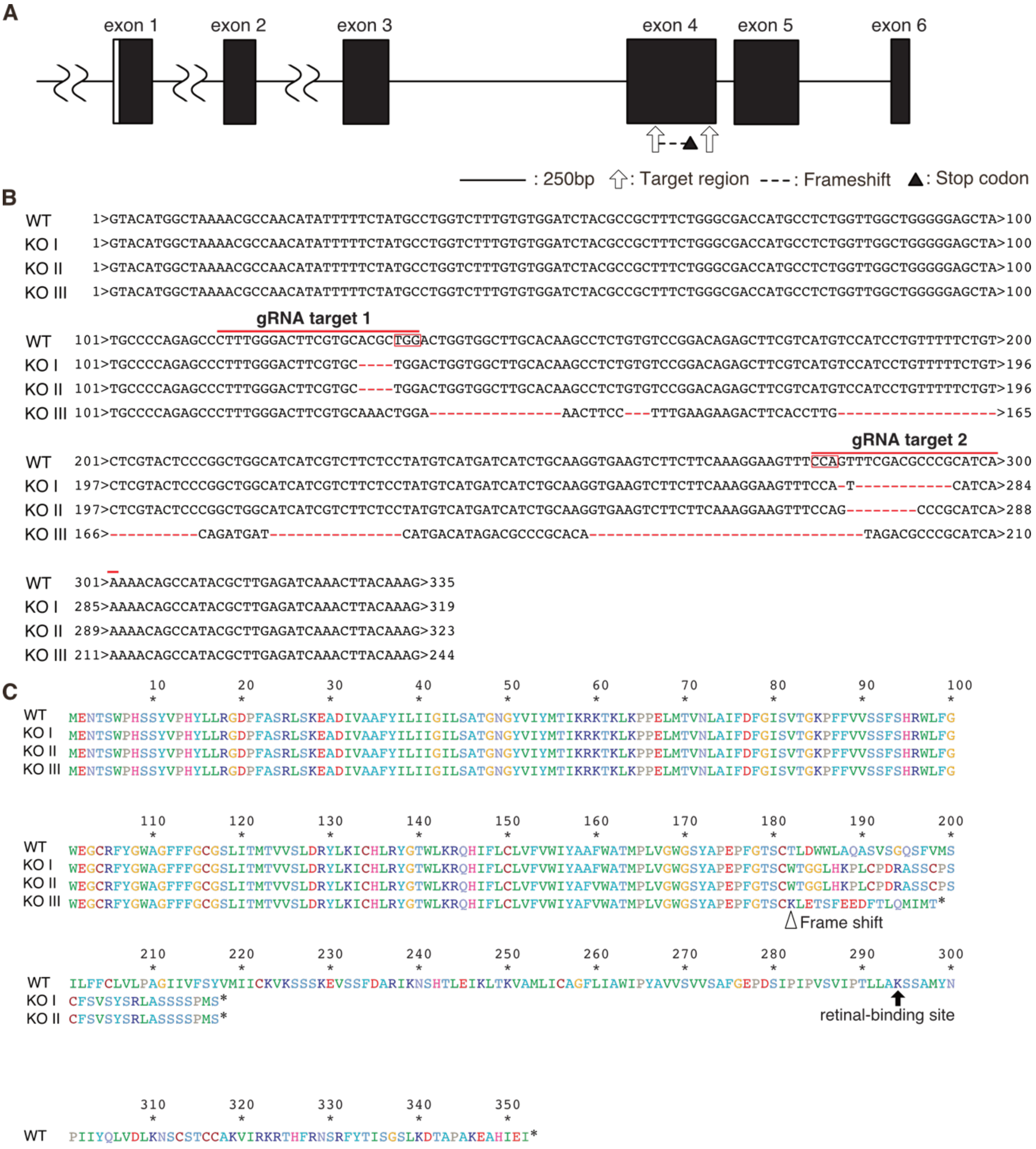


**Fig. S9.** ***opn5m*** **knockout medaka generated in this study.**

**(A)** Intron-exon structure of *opn5m* genes is illustrated based on ENSORLT00000019589.2. Target regions of CRISPR/Cas9 are indicated by the arrow. **(B)** Alignment of genomic DNA sequence of knockout medaka in exon 4. We generated and used three patterns of mutation. KO I: 4 base pair (bp) and 12 bp deletion, KO II: 4 bp and 8 bp deletion, KO III: 149 bp deletion and 57 bp insertion. gRNA targets of the CRIPSR are underlined, and PAM sequences are shown in squares. **(C)** Alignment of deduced amino acid sequences of wild type (WT) and knockout of *opn5m* gene (KO I, II, III). These knockout genotypes of *opn5m* have lost the K294, the retinalbinding site indicated by black arrow, suggesting the null/loss of function. The start position of the frameshift mutation in the amino acid sequences of Opn5m is indicated by white arrowheads. *; stop codon.

**Fig. S10**

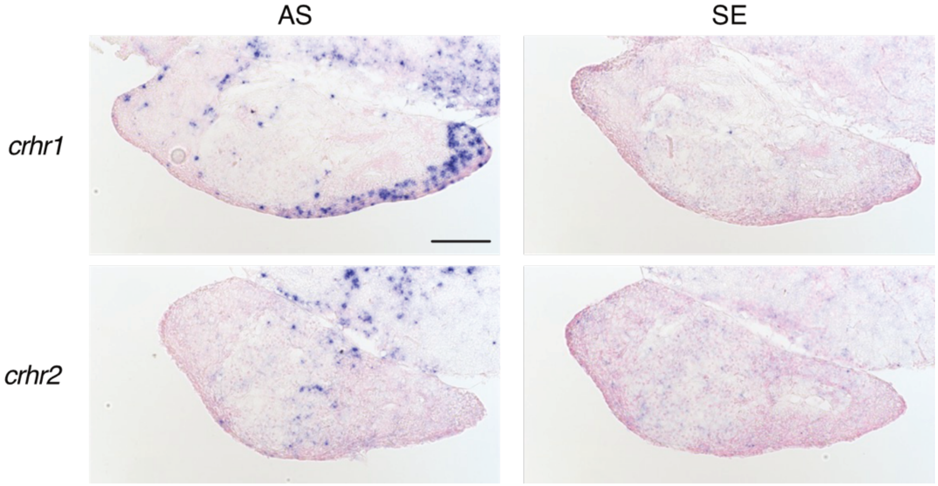


**Fig. S10. *In situ* hybridization suggests that *crhr1* is expressed in *pomc* cells in the pituitary.**

Expression of CRH receptor (*crhr1*, *crhr2*) in the pituitary analyzed by *in situ* hybridization. The photographs indicate that *crhr1* is localized in *pomc*-expressing regions. Scale bar, 100 µm.

**Fig. S11**

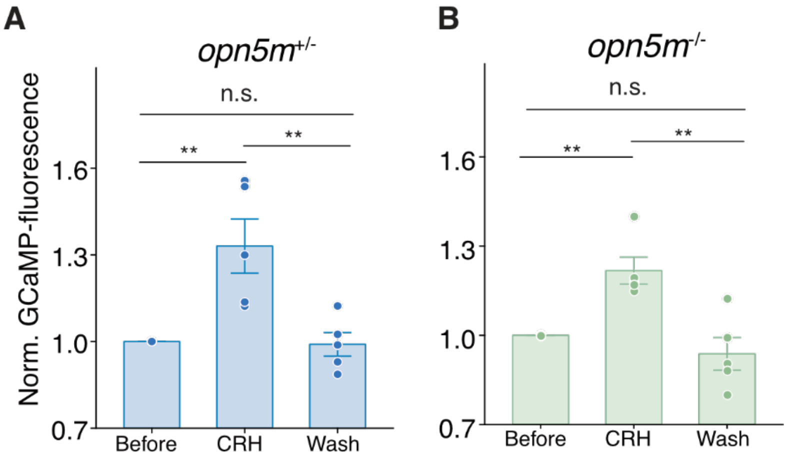


**Fig. S11. Both *opn5m* $^{+/-}$ and *opn5m* $^{-/-}$ medaka pituitary respond to CRH.**

Application of CRH significantly increases the GCaMP fluorescence in melanotrophs and this increase is abolished by washout. The effect of CRH is similarly observed in both in *opn5m*$^{+/-}$ and *opn5m*$^{-/-}$ medaka. Before; before CRH perfusion, CRH; during 400 nM CRH perfusion, Wash; washout after CRH perfusion. $n$ = 5 medaka, **; $P$ = 0.005, $P$ = 0.004, $P$ = 0.007, $P$ = 0.001, respectively; n.s., not significant; Tukey test.

**Fig. S12**

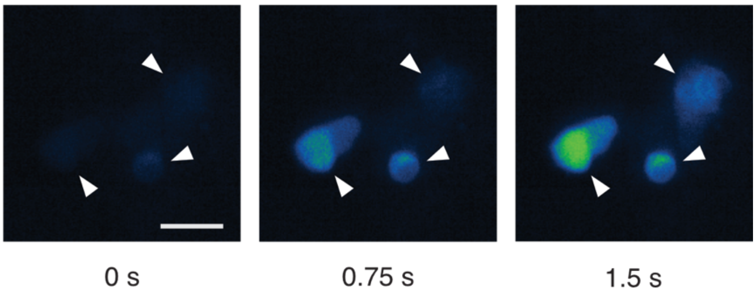


**Fig. S12. LβT2 cells transfected with *opn5m* and *gcamp6s* also respond to the blue excitation light.**

Similar to HEK293A cells, LβT2 cells that express Opn5m and GCaMP6s (indicated by white arrowheads) show $[Ca^{2+}]_i$ increase in response to the excitation light. The graph shows the time course of GCaMP fluorescence images of LβT2 cells. $n$ = 71 cells. Scale bar, 25 µm. Data are represented as mean ± SEM.

**Fig. S13**

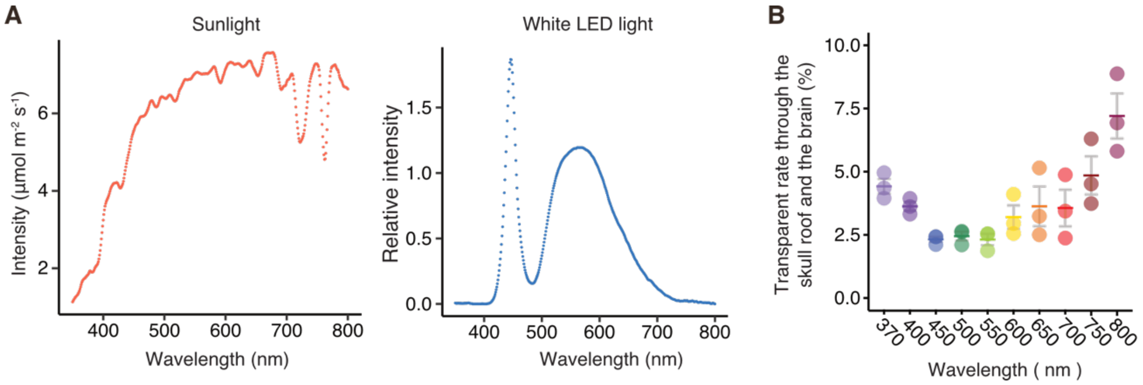


**Fig. S13. Natural sunlight from outside the body should induce the $[Ca^{2+}]_i$ response of melanotrophs observed in the present study.**

**(A)** To estimate the amount of light that reaches the pituitary in adult medaka in the natural state, we examined the transparency of light of various wavelengths through the entire structure dorsal to the pituitary, including the skull and brain. This graph shows the rate of various light transparency. Approximately 2 to 4% natural sunlight was estimated to reach the pituitary in adult wild strain medaka. $n$ = 3 medaka. **(B)** The spectrum of the sunlight and white LED light we used in the experiments in Fig. 3C to D. The amplitude of light activation on Opn5m of medaka pituitary that naturally occurs is calculated as follows. Considering the absorption spectrum of 11-*cis*-retinal-bound Opn5m (= y0 + A*exp (-(((x-x0)/width) ^2)), y0 = 0, A = 0.6593, x0 = 363.28, width = 52.064), transparency of the structure above the pituitary, and the spectrum of sunlight and white LED, ~2667 µmol $m^{-2}$ $s^{-1}$ sunlight (sunny day, July 27, 2023, around noon, Kashiwa, Chiba, Japan) is equivalent to 255 µmol $m^{-2}$ $s^{-1}$ white LED. Thus, the $Ca^{2+}$ response induced by the white LED light up to 255 µmol $m^{-2}$ $s^{-1}$ should occur under the natural sunlight, *in vivo*.

**Fig. S14**

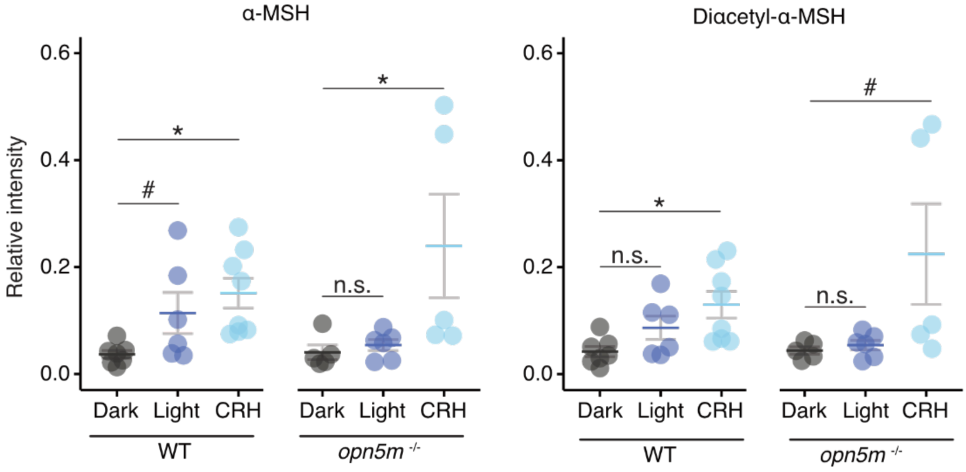


**Fig. S14. α-MSH peptide showed a trend of release by the light.**

LC-MS results showing MSH derivatives other than desacetyl α-MSH are shown in the main figure. α-MSH tended to be released by light in wild-type medaka. In contrast, α-MSH release is not observed in response to the light in *opn5m*$^{-/-}$ medaka pituitary. $n$ = 5–8, #, $P$ = 0.12, $P$ = 0.055; *, $P$ = 0.011, $P$ = 0.042, $P$ = 0.011, respectively; n.s., not significant; Dunnet test. Data are represented as mean ± SEM.

**Fig. S15**

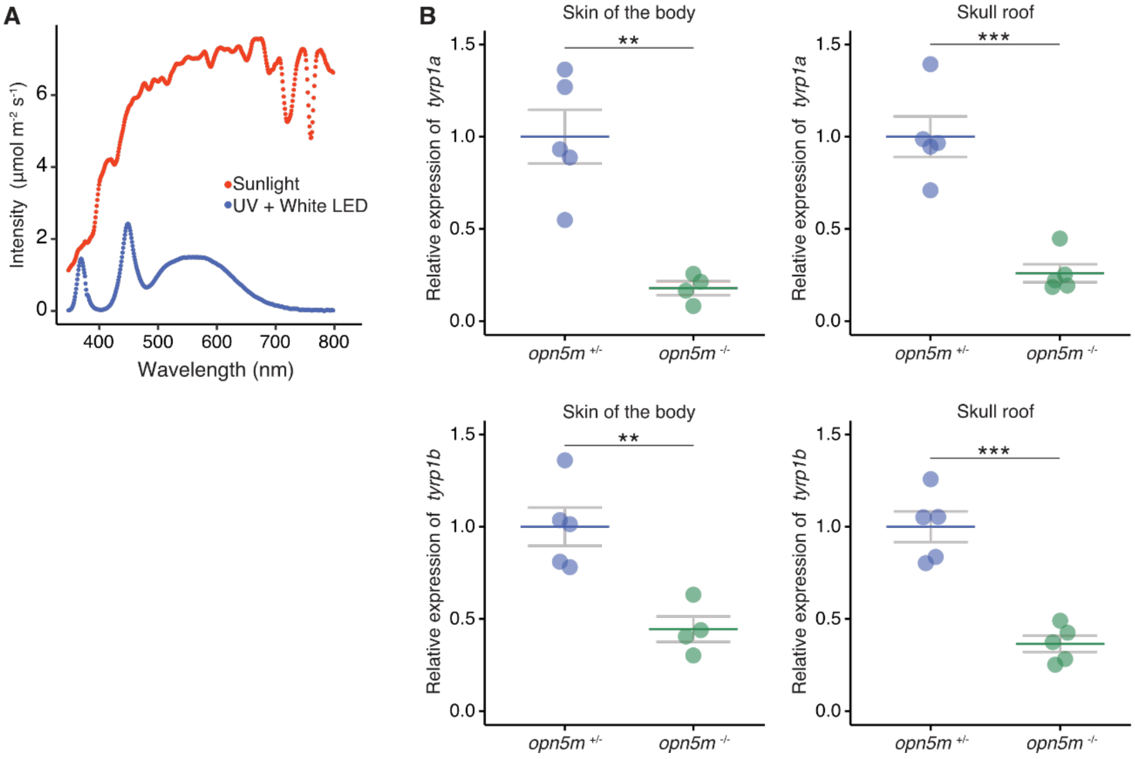


**Fig. S15. The expression level of tyrosinase-related protein in the skin and skull was lower in *opn5m*$^{-/-}$ medaka than in *opn5m*$^{+/-}$ medaka.**

**(A)** The irradiation light spectrum of the LED setup used in the *in vivo* experiment and the sunlight. Based on these spectra and the absorption spectrum of Opn5m, we estimated the relative effect of the LED setup used in the *in vivo* experiment. The result of the calculation detailed in the material and methods section indicated that the LED light used in the *in vivo* experiment was less than one-quarter compared to the sunlight in terms of Opn5m activation (~21%). **(B)** The relative expression level of tyrosinase-related protein 1a (*tyrp1a*) or 1b (*tryp1b*) in the skin of the body or the skull roof of *opn5m*$^{+/-}$ or *opn5m*$^{-/-}$ medaka. The expression of both tyrosinase-related protein genes was suppressed in *opn5m*$^{-/-}$ medaka. $n$ = 5 or 4. **, $P$ = 0.0018, $P$ = 0.004; ***, $P$ = 0.00028, $P$ = 0.00014, respectively; Student's $t$ test. Data are represented as mean ± SEM.

**Table S1. List of primer sequences.**

| Primer | Sequences (5' to 3') |
|---|---|
| *pomc* UP3.7k_SE (backbone link) | ATGCTTGCTCACATGTGCAACAGACTGGGATAGGCTCC |
| *pomc* UP_AS (GCaMP link) | GTCGACCATGGTGGCCCTCTGTGAAGCAAAAAAAAACACAATC |
| **Single *in situ* hybridization** | |
| *pomc* SE | AGAAGAGGATCAAGGGAAATCTTCA |
| *pomc* AS | TCATTGCTTGCAAATAAATCTCTTT |
| *crhr1* SE | TCAGAGCGATGTGGACTGCTTGAT |
| *crhr1* AS | GACAGTTCATACCTGACCGGTGGTG |
| *crhr2* SE | CTCTCCGCTTGTCGTTTAGCCTCG |
| *crhr2* AS | TGGAAAAGCAAAGGTTTTTGTCGATGT |
| **Genotyping** | |
| *opn5m* SE1 | CCTTCAATCTTAGGTACATGGCTAA |
| *opn5m* AS1 | TGATACTTTTAGTCTAAGTCTGTGT |
| *opn5m* SE2 | AGTAGTGGCTGTGTGACATGG |
| *opn5m* AS2 | GCACAAATCAGCATTGCCACCT |
| **qPCR** | |
| *actb* SE | GTGATGTTGATATCCGTAAGGATCTGTA |
| *actb* AS | TCTGGTGGGGCAATGATCTTGA |
| *tyrosinase* SE | TCTGTCTTCTCCTCATGGAAGGTAATC |
| *tyrosinase* AS | GGTTACGCAACAGTGGACCTT |
| *tyrp1a* SE | GAGCGCGACATGCAGGACAT |
| *tyrp1a* AS | GAGTTGGAGTCAAAAGTGCTC |
| *tyrp1b* SE | GATCAGAGAGTCTCTGTGGTAAG |
| *tyrp1b* AS | GGCCCAACACCTCCTGATAC |

**Table S2. List of probe sequences.** All Oligos was used according to the Kishi *et al*., 2019 (*37*).

| Oligo name | Sequence | Usage |
|---|---|---|
| Clean.G | CCCCGAAAGTGGCCTCGGGCCTTTTGGCCCGAGGCCACTTTCG | Pre-incubate before primer addition to remove contaminating guanine |
| h.28.28.tail | ACAACTTAACGGGCCTTTTGGCCCGTTAAGTTGTGTTAAGTTGTTTTTTT | Extension of probe or branch oligos |
| h.31.31.tail | ATTATTCACTGGGCCTTTTGGCCCAGTGAATAATAGTGAATAATTTTTTT | Extension of probe or branch oligos |
| h.73.73.tail | ATTCCTAATCGGGCCTTTTGGCCCGATTAGGAATGATTAGGAATTTTTTT | Extension of probe or branch oligos |
| h.27.27.tail | ACATCATCATGGGCCTTTTGGCCCATGATGATGTATGATGATGTTTTTTT | Extension of probe or branch oligos |
| h.28.31.tail | ATTATTCACTGGGCCTTTTGGCCCAGTGAATAATGTTAAGTTGTTTTTTT | Remap primer sequence in probe from no.28 to no.31 |
| h.28.73.tail | ATTCCTAATCGGGCCTTTTGGCCCGATTAGGAATGTTAAGTTGTTTTTTT | Remap primer sequence in probe from no.28 to no.73 |
| 31*.31*.31*.31*.27 | AGTGAATAATAGTGAATAATAGTGAATAATAGTGAATAATTTCATCATCAT | Branch for signal amplification in SABER |
| 27*.27*.27*.28 | ATGATGATGTATGATGATGTATGATGATGTTTCAACTTAAC | Branch for signal amplification in SABER |
| 28*.488 | /5ATTO488N/ttGTTAAGTTGtGTTAAGTTGt | Fluorescent detection |
| 73*.565 | /5ATTO565N/ttGATTAGGAAtGATTAGGAAt | Fluorescent detection |

| Remapping information in PER reaction |
|---|
| Primer sequence of OlOpn5m.28 was remapped to no.31. |
| Primer sequence of Olpomca.28 was remapped to no.73. |
| Primer sequences of brances for amplification were not remapped. |

**Applied probes and branches in each hybridization step**

| Tissue | probe hybridization | 1st branch hybridization | 2nd branch hybridization | Fluorescent detection |
|---|---|---|---|---|

| | | | | |
|---|---|---|---|---|
| medaka pituitary | OlOpn5m.31 (extended probe) | 31*.31*.31*.31*.27 (extended branch) | 27*.27*.27*.28 (extended branch) | 28*.488 |
| | Olpomca.73 (extended probe) | N.A. | N.A. | 73*.565 |

| **Oligo pool name** | **Sequence** |
|---|---|
| OlOpn5m.28 | TAAAGTCAGCAGTGTGGCAGGTAGGGGTGTGCAAAAtttCAACTTAAC |
| OlOpn5m.28 | AACTTCACGAGTCCAAAGCTTTCATGGTCGGTTCTAtttCAACTTAAC |
| OlOpn5m.28 | GGGAACGTTGCAGAAATAGAAAACAGTGAAACGTCATGAGAtttCAACTTAAC |
| OlOpn5m.28 | ATGTGTTCTCCATTACTGTCGTCAGTGTGGAAGGTCtttCAACTTAAC |
| OlOpn5m.28 | GGAGATAATGCGGGACATACGAAGAGTGAGGCCACGtttCAACTTAAC |
| OlOpn5m.28 | AATGACGTAGCCATTTCCTGTTGCAGACAGGATTCCtttCAACTTAAC |
| OlOpn5m.28 | AGGCTTCAGCTTCGTCTTGCGTTTGATGGTCATGTAtttCAACTTAAC |
| OlOpn5m.28 | ACAGCCAGCGGTGGGAGAAACTAGACACAACGAAAAtttCAACTTAAC |
| OlOpn5m.28 | GTCCAGGCTCACCACTGTCATTGTGATGAGGCTCCCtttCAACTTAAC |
| OlOpn5m.28 | GCGGCGTAGATCCACACAAAGACCAGGCATAGAAAAtttCAACTTAAC |
| OlOpn5m.28 | ACATGACGAAGCTCTGTCCGGACACAGAGGCTTGTGtttCAACTTAAC |
| OlOpn5m.28 | TGACATAGGAGAAGACGATGATGCCAGCCGGGAGTAtttCAACTTAAC |
| OlOpn5m.28 | GGAAACTTCCTTTGAAGAAGACTTCACCTTGCAGATGATCAtttCAACTTAAC |
| OlOpn5m.28 | AAGCTATCAAGAAGCCTGCACAAATCAGCATTGCCAtttCAACTTAAC |
| OlOpn5m.28 | ACGCTGAAACCACTGAGACCACTGCATATGGAATCCtttCAACTTAAC |
| OlOpn5m.28 | ACAGATACAGGAATGGGAATGGAATCTGGTTCACCAAtttCAACTTAAC |
| OlOpn5m.28 | TGTACATCGCTGAAGACTTAGCCAACAATGTAGGGATGtttCAACTTAAC |
| OlOpn5m.28 | TCCTTATGACTTTAGCACAGCAGGTTGAGCATGAGTTTTtttCAACTTAAC |
| OlOpn5m.28 | ATTTCAATGTGAGCTTCTTTGGCGGGCGCTGTGTCCtttCAACTTAAC |
| OlOpn5m.28 | CAGGGAACAAGGCATACTGTCCAAACCGAAAGAGGCtttCAACTTAAC |
| OlOpn5m.28 | GCAGCGTGTGATTGGATAGGTGTCGAGAACAGAGGGtttCAACTTAAC |
| OlOpn5m.28 | AGCAACAACGGTGAGCTGTTAATGATTTCCCAGCAGtttCAACTTAAC |
| OlOpn5m.28 | TGAGGTGGCGAAATCAAGCTGGAAGACTCAGGAACCtttCAACTTAAC |
| OlOpn5m.28 | CCTTTTGCTTTAGCAATGGTCAGACAGGTTCTCTGTAACATtttCAACTTAAC |
| OlOpn5m.28 | TCCAGTTTCAAAACCTGAGTTATGGCTTTCATCTCAAACGTtttCAACTTAAC |
| OlOpn5m.28 | ACAACATCCTGTACTAGCTGCATAAATGCAACTCAAGAGTtttCAACTTAAC |
| OlOpn5m.28 | TGTGACCATTGAAGAAGTGTCTTCTTAGCCATTTATCGGTTtttCAACTTAAC |
| OlOpn5m.28 | CCGAACCTTGAACTGTTGCCTTACGATCATTGACGAATAAAtttCAACTTAAC |
| OlOpn5m.28 | GGAGCAGGCCTTATTAAGACTTTACAGCAGGACAGTTTTtttCAACTTAAC |

| | |
|---|---|
| OlOpn5m.28 | CCTGTCGGTCAAAGCAATCTTCAGGGTCAGTGAAAAtttCAACTTAAC |
| OlOpn5m.28 | AGATGTAAAGTCACCCACGTTGGCTTCGATAATCTTCTTtttCAACTTAAC |
| OlOpn5m.28 | TCAGAGCCATTTGAACTAAATTTGTCCCTGAAAGAGTCGTTtttCAACTTAAC |
| OlOpn5m.28 | GTGGATAATGAGTAGCCTCTAAATGTTGGGAAGCAACAAGAtttCAACTTAAC |
| OlOpn5m.28 | TTAGGGATAACGCCCCGGGCCTTTCCAACATGTAGGtttCAACTTAAC |
| OlOpn5m.28 | TGGTGACTGTACCCAGTCAGGAATAGTTACATAGTTTGAGTtttCAACTTAAC |
| OlOpn5m.28 | GTGGTAACTAAGATAAGCAACGAGAACCACTGTATTGCAGTtttCAACTTAAC |
| OlOpn5m.28 | GGTGTGCACCTAGAACATTTTGCCAGTCCACTGCAGtttCAACTTAAC |
| OlOpn5m.28 | GGCTTGGAACAGATGGTGTGATGGTAGTTAACCACATTTTAtttCAACTTAAC |
| OlOpn5m.28 | CCTTTCTTGAATGACTCCCTGGGATGCTTAACTTTACCAAAtttCAACTTAAC |
| OlOpn5m.28 | CCATCTTCAAAATGTTGAGCAAAGGCAGATTTCCCAGAAAGtttCAACTTAAC |
| OlOpn5m.28 | CCAACAGCACTAGTGTCCATGTATCCTGAAACATCTAAGTCtttCAACTTAAC |
| OlOpn5m.28 | ACAGTGTGGCGTATTTCACAGATTTTACATCTACCAAACCAtttCAACTTAAC |
| OlOpn5m.28 | CACAGCTGTTCTTTTGAGCAAACAGGAAATCTCTCCTCtttCAACTTAAC |
| OlOpn5m.28 | TGACTGATTTTCTTTCCACTCAACCCAGTGTCCTTCAAtttCAACTTAAC |
| OlOpn5m.28 | GCGACCTTTGATTGACCCCAATGTTGGAAGAAGTTTCTtttCAACTTAAC |
| OlOpn5m.28 | CCGCATGTGGGAGAACTTAATTAACACTGTCACAAGTAGAGtttCAACTTAAC |
| OlOpn5m.28 | GTCCAATTCTGATGTCGAATTATTCCAGGTCCTTGACACATtttCAACTTAAC |
| OlOpn5m.28 | TCTGTGGATTCGAGAAGTTTAGAAATGAGGCTGCAATGTAAtttCAACTTAAC |
| OlOpn5m.28 | TCCAGTGTTCAGAGGAGATATGGCTTTGATCAAACCTAAGAtttCAACTTAAC |

| **Oligo pool name** | **Sequence** |
|---|---|
| Olpomca.28 | AGCAGAGTTGGACTTTTATATTCTGCTGACCAAGGACGtttCAACTTAAC |
| Olpomca.28 | AGATTTCCCTTGATCCTCTTCTGCTCTGATGCTTCGtttCAACTTAAC |
| Olpomca.28 | AAAGAGAAGGGATCTGAGGGAGGTGGAGGCTGGAGGtttCAACTTAAC |
| Olpomca.28 | GAGTAGGAGCGTTTGGTTTGAGGGGATGGAGAGATGtttCAACTTAAC |
| Olpomca.28 | TCCTCCTCCACTCCGTTTGGGGTGTAGACCTTGACGtttCAACTTAAC |
| Olpomca.28 | CGCCTCCTCATCTCACCTGGGAAAACCTCAGAGGACtttCAACTTAAC |
| Olpomca.28 | CTTGTAGGAGCCATCCTTCTTCTCCTGCAGGCCAGCtttCAACTTAAC |
| Olpomca.28 | TCTTCATGAAGCCGCCATAGCGTTTACTGGCGGGAGtttCAACTTAAC |
| Olpomca.28 | GAAGAGCGTCACCAGTGGTTTCTGGCGATCCTCCTCtttCAACTTAAC |
| Olpomca.28 | CAAAAGAACCAATGAGAACTCTTTGCTGCCTGCTGATTATTtttCAACTTAAC |
| Olpomca.28 | CGCCAAAACACTGCAGTTTCAGAGTCTTTTATTGGGAAtttCAACTTAAC |
| Olpomca.28 | ATGTGCTATAGGGTGCACCTTATGATCTGGTGCCCCtttCAACTTAAC |
| Olpomca.28 | GATAGGGATGCCATTTCCGTTCATGTGCGAGTCAAAtttCAACTTAAC |
| Olpomca.28 | AGTCTGGACCTAAACCTGAAAGAACACCTTAGGGATGAATTtttCAACTTAAC |

| | |
|---|---|
| Olpomca.28 | GAAGTGGCCAGTTCTCTGGTCATTGTTCTCGCTGATtttCAACTTAAC |
| Olpomca.28 | GGCAAGGGCTGATGTGAACTGTAGTATGCAAACCAGtttCAACTTAAC |
| Olpomca.28 | GCGACACAATCGAGTAAACCCAGGTCATGAATATAAGGTTtttCAACTTAAC |
| Olpomca.28 | GAGGGACAAGGTGTGACTGCTGTCCAGTGACGATAAtttCAACTTAAC |
| Olpomca.28 | ACTCTGGTAGTCATCTGTTTCGGGTTTGGATTTGGAAAATtttCAACTTAAC |
| Olpomca.28 | TCAGATTGTTTCTAACAGACTGCACCGCAAGTTCGTtttCAACTTAAC |
| Olpomca.28 | AGGAGCTCAGTTGAAGCTTAAGATATACGTCCAGTCTGGtttCAACTTAAC |
| Olpomca.28 | GGAACAGAGCATCTAGAGGTCCTGAATTCAAGATCAGTGTTtttCAACTTAAC |
| Olpomca.28 | AAACTGGTCCCCTTATTGTCCACGCGCACAGAGTGTtttCAACTTAAC |
| Olpomca.28 | GCTCTTATGCCACACTTCACTCATTAGTAAGGGGCAAtttCAACTTAAC |
| Olpomca.28 | TCAAAAGCCATAAACTGGAAGCCAATTCCTGCAGTGtttCAACTTAAC |
| Olpomca.28 | GCGCCGGGGTTCACATCCTATCTTATAATAAACAAAGACTTtttCAACTTAAC |
| Olpomca.28 | GTAGCAGACTTTACGGACAGCCCCACCCCAATGTCCtttCAACTTAAC |
| Olpomca.28 | CGCTCCTTCAGTGTGATCGAAACATTTGTGGTTGAAGtttCAACTTAAC |
| Olpomca.28 | TCTGTGTGTATTAGCCAAACATTCCACTTCAACATGGAGTTtttCAACTTAAC |
| Olpomca.28 | CGCTCCGTACCCATCTATGAAATGAAGTTTCCAGACCtttCAACTTAAC |
| Olpomca.28 | CGTCCTCCTCCGTCTCCATCTGGACTTTCTCCTCCTtttCAACTTAAC |
| Olpomca.28 | CGCATTGTAGCAGAAGTTGATCCAGATCAAGCATTATCCAAtttCAACTTAAC |
| Olpomca.28 | TGGTATCGAAACTACTTATACTAAGATCAATCCGCCCACCCtttCAACTTAAC |
| Olpomca.28 | CGCGCATGAGTTAACTCGAGCTCTTTACTGTATGTGTTAAAtttCAACTTAAC |

### References


31. T. Ishikawa *et al.*, Medaka as a model teleost: characteristics and approaches of genetic modification. *Laboratory Fish in Biomedical Research*, (2022).
32. K. Maruyama, B. Wang, Y. Ishikawa, S. Yasumasu, I. Iuchi, 1kbp 5′ upstream sequence enables developmental stage-specific expressions of globin genes in the fish, medaka Oryzias latipes. *Gene* **492**, 212-219 (2012).
33. B. Zempo, S. Kanda, K. Okubo, Y. Akazome, Y. Oka, Anatomical distribution of sex steroid hormone receptors in the brain of female medaka. *Journal of Comparative Neurology* **521**, 1760-1780 (2013).
34. Y. Kawabata-Sakata, Y. Nishiike, T. Fleming, Y. Kikuchi, K. Okubo, Androgen-dependent sexual dimorphism in pituitary tryptophan hydroxylase expression: relevance to sex differences in pituitary hormones. *Proc Biol Sci* **287**, 20200713 (2020).
35. A. Edelstein, N. Amodaj, K. Hoover, R. Vale, N. Stuurman, Computer control of microscopes using μManager. *Curr Protoc Mol Biol* **Chapter 14**, Unit14.20 (2010).

36. B. J. Beliveau *et al.*, OligoMiner provides a rapid, flexible environment for the design of genome-scale oligonucleotide in situ hybridization probes. *Proceedings of the National Academy of Sciences* **115**, E2183-E2192 (2018).
37. J. Y. Kishi *et al.*, SABER amplifies FISH: enhanced multiplexed imaging of RNA and DNA in cells and tissues. *Nature Methods* **16**, 533-544 (2019).
38. R. Schmid *et al.*, Integrative analysis of multimodal mass spectrometry data in MZmine 3. *Nature Biotechnology* **41**, 447-449 (2023).
39. R_Core_Team*,* R. F. f. S. Computing, Ed. (2022).
40. H. Ando, K. Ukena, S. Nagata, *Handbook of Hormones: Comparative Endocrinology for Basic and Clinical Research*. (Elsevier Science, 2021), pp. 211.
41. T. D. Lamb, Photoreceptor spectral sensitivities: Common shape in the long-wavelength region. *Vision Research* **35**, 3083-3091 (1995).
42. V. I. Govardovskii, N. Fyhrquist, T. O. M. Reuter, D. G. Kuzmin, K. Donner, In search of the visual pigment template. *Visual Neuroscience* **17**, 509-528 (2000).
43. T. Yamashita *et al.*, Opn5 is a UV-sensitive bistable pigment that couples with Gi subtype of G protein. *Proceedings of the National Academy of Sciences* **107**, 22084-22089 (2010).
